# BCL6 in T cells promotes type 1 diabetes by redirecting fates of insulin-autoreactive B lymphocytes

**DOI:** 10.1101/2025.08.25.671997

**Authors:** Landon M. Clark, Jack C. McAninch, Dudley H. McNitt, Marguerite L. Padgett, Tyler W. Jenkins, Lindsay E. Bass, Casey M. Nichols, Jeffrey C. Rathmell, Rachel H. Bonami

## Abstract

Currently approved type 1 diabetes (T1D) immunotherapies broadly target T cells and delay but do not fully prevent diabetes development, highlighting the need for more selective targets. Anti-insulin germinal center B cells are uniquely able to present pathogenic insulin epitopes and drive anti-insulin T cells to adopt a T follicular helper fate. T cell expression of BCL6, a key transcriptional repressor in the germinal center response, is essential for spontaneous diabetes in non-obese diabetic (NOD) mice. However, the impact of T cells on pro-pathogenic anti-insulin B cell activity is still poorly understood. Here, we show that VH125^SD^.NOD mice with T cell loss of BCL6 still produce peripheral anti-insulin B cells yet are protected against diabetes (relative to *Bcl6*-sufficient controls). This protection was associated with reduced activation, proliferation, germinal center differentiation, and pancreatic infiltration of insulin-binding B cells. Minimally supervised analysis revealed insulin-binding B cells skew towards atypical memory B cell subsets specifically in pancreas and pancreatic lymph nodes, which was reduced by *Bcl6*^Δ*CD4*^ loss. Overall, this work suggests BCL6-expressing T cells are pivotal to license pathogenic insulin-binding B cells. Our findings support BCL6 inhibition as a promising T1D immunotherapy, even after insulin autoimmunity is established in the B cell repertoire.

**Article Highlights:**

- Loss of floxed *Bcl6* via *Cd4*-Cre protects against type 1 diabetes even when an insulin-skewed B cell repertoire is present
- BCL6 loss in T cells reduces anti-insulin B cell upregulation of T cell co-stimulatory molecules, proliferation, and IgG class switching in pancreas and pancreatic lymph nodes in VH125^SD^.NOD mice
- Anti-insulin B cells skew towards atypical and atypical memory B cell phenotypes compared to non-insulin binding B cells in pancreas and pancreatic lymph nodes, only some of which are reduced by T cell loss of *Bcl6*
- This study highlights the translational potential of targeting BCL6, even after the establishment of insulin-reactive B cells, in line with typical intervention points for at-risk individuals

**Graphical Abstract:** 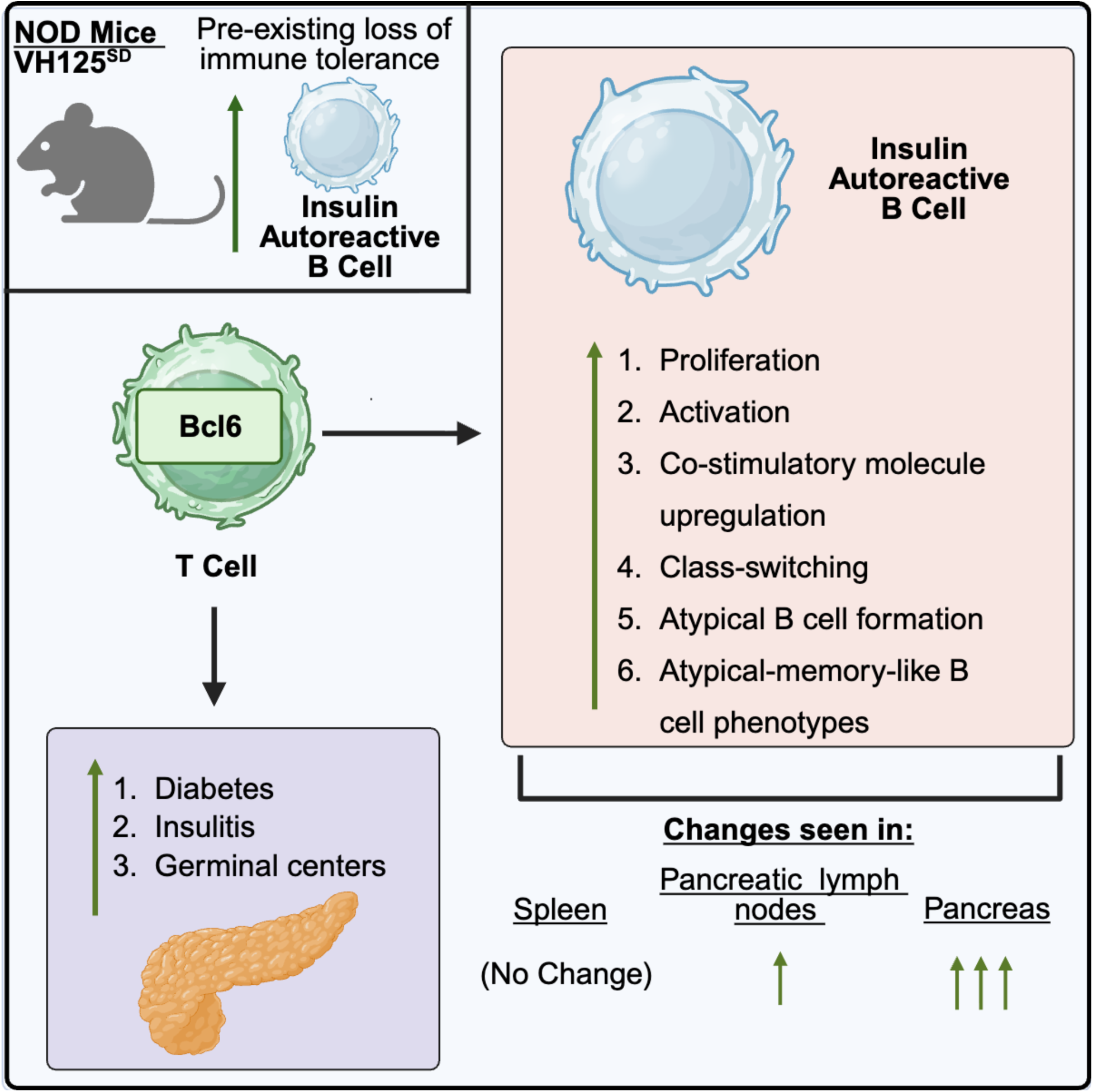

## Introduction

Immunotherapies targeting T or B cells are a mainstay treatment in rheumatic disease management and are beginning to be applied to type 1 diabetes (T1D) treatment (e.g. teplizumab), but no current treatment durably halts beta cell demise ^1–3^. Rituximab trials for T1D management showed reduced beta cell decline but did not significantly delay disease progression, highlighting the urgent need to identify new T1D immunotherapies^4^. A major challenge in the clinical management of many autoimmune diseases is that individuals typically present for disease diagnosis well after initial immune tolerance breach for autoantigens has occurred, as demonstrated by the predictive power of autoantibodies in diseases such as SLE, RA, and T1D ^5–7^. Thus, new T1D immunotherapies should ideally reduce disease burden even when initiated after autoreactive B cell tolerance breach has occurred (signaled by autoantibody seropositivity) ^8,9^.

Insulin is a major autoantigen in T1D; insulin autoantibodies predict T1D in mice and humans and are associated with earlier age of T1D onset ^10^. Anti-insulin B cells can act as antigen-presenting cells to drive CD4+ T cell activation ^11^. Recent evidence suggests one particular CD4+ T cell subset, T follicular helper cells (Tfh) cells, as being involved in T1D. Tfh cells are important for the germinal center (GC) response, where B cells can interact with Tfh cells to support B cell affinity maturation and generation of memory B cell and long-lived plasma cell responses ^12,13^. Circulating Tfh cells are elevated in T1D patients and abatacept response heterogeneity is linked to differences in Tfh-like populations, implicating Tfh cells in T1D development ^14–18^.

The role of GCs in T1D is still unclear, with data supporting and refuting their necessity in driving disease. In support of GCs driving T1D pathology, anti-insulin (VH125^SD^) B cells with a GC, but not non-GC phenotype, drive anti-insulin (8F10) T cells to undergo activation and proliferation *in vitro* ^19^. This was attributed to unique antigen processing/presentation capacity acquired by GC B cells, which enabled them to generate the insulin B:12-20 epitope recognized by 8F10 T cells in mice and pathogenic T cell clones in humans ^19,20^. Anti-insulin (VH125^SD^) B cells and anti-insulin (8F10) T cells drive each other to differentiate into GC B cells and Tfh cells, respectively, and elicit insulin autoantibody production (otherwise silenced in the VH125^SD^ T1D-prone non-obese diabetic (NOD) model) ^19^. Thus, anti-insulin B and T cells influence one another to defy immune tolerance mechanisms that otherwise restrain their differentiation capacity ^19^.

Evidence against T1D dependence on GCs also exists. Loss of SLAM-associated protein (SAP) in NOD mice led to a near complete loss of GCs, yet resulted in only modest diabetes protection, with ∼50% of mice still developing diabetes ^21^. The unexpected persistence of a population with a Tfh phenotype (defined as CXCR5^hi^ PD1^hi^ BCL6^+^ ICOS^hi^ CD44^hi^) in *SAP*-deficient NOD mice may help explain the incomplete disease protection observed in this model. This calls attention to the possibility for altered molecular governance of pathogenic Tfh differentiation and maturation in NOD mice, relative to Tfh that govern other protective/autoimmune settings ^21,22^.

To better understand the role of GCs in NOD mice, we recently generated *Cd4-*Cre- mediated deletion of the transcriptional repressor, BCL6, in NOD mice ^23–26^. We showed that T cell loss of BCL6 abolishes Tfh and GC B cells but provided complete protection against diabetes ^27^. However, this mechanism is still poorly understood, as *Cd4-*Cre eliminates *Bcl6* in both CD4 and CD8 T cell populations due to the double-positive stage in the thymus ^28^. While other non-T cell populations can also express CD4 (e.g., CD4+ dendritic cells, ^29^), we will refer to this model as T cell-specific loss of *Bcl6* hereafter but acknowledge the potential role of other CD4+ populations. Additionally, how BCL6 loss in T cells impacts autoreactive B cells is unclear. Insulin-binding B cells selectively upregulate CD86 in the pancreas relative to non-insulin-binding B cells ^30^, but it is unclear whether this and other pro-pathogenic changes in insulin-binding B cells depends on BCL6-expressing T cells.

While activated B cells can enter the GC response, they can also enter an extrafollicular (EF) route, either independently of the GC response or through premature exit of the GC, to form atypical (or age-associated) B cells and short-lived plasmablasts ^31–33^. Tfh cells support EF responses by enhancing atypical B cell generation, which accumulate in the settings of aging, autoimmunity, and some infection models ^34–38^. Atypical B cells are heterogeneous, with different subpopulations marked by CD11c, CD11b, and/or T- bet expression ^36^. T-bet+ CD11c+ B cells that arise in infection depend on BCL6-expressing Tfh cells but are thought to chiefly arise independently of the GC ^37^. Atypical B cells have not been well characterized in T1D but are implicated in other autoimmune diseases such as SLE ^39,40^. Thus, these populations were examined here.

Given that autoreactive B cell populations are rare and difficult to study in NOD mice, we deploy the VH125^SD^.NOD model, whereby an IgH locus-targeted B cell receptor (BCR) heavy chain transgene enables a small (1-3%) but reliably trackable population of insulin autoreactive B cells to form in the bone marrow ^41^. Here, we show that despite VH125^SD^ BCR transgene-forced expansion of anti-insulin B cells, T cell loss of *Bcl6* led to nearly complete prevention of diabetes in VH125^SD^ mice. Elimination of BCL6 from T cells led to a reduction in anti-insulin B cell expression of T cell costimulatory molecules, spontaneous proliferation *in vivo*, IgG switching, and pancreatic islet infiltration by T and B lymphocytes. We observed anti-insulin B cell skewing towards atypical B cell subsets and a CD80+ PDL2+ T-bet+ subset that could represent atypical memory, particularly in the pancreatic lymph nodes and pancreas, compared to non-insulin-binding B cells.

Some, but not all, of these skewed populations were lost with *Cd4*-Cre deletion of *Bcl6*. These results highlight the importance of BCL6 expression, possibly in CD4 and/or CD8 T cells, to activate and license anti-insulin B lymphocytes as APCs and support beta cell attack in T1D.

## Results

### BCL6 in T cells promotes spontaneous GC formation, diabetes, and anti-insulin B cell infiltration of pancreatic islets in VH125^SD^.NOD mice

*Cd4*-Cre *Bcl6* loss prevents T1D in NOD mice ^27^, but it is currently unclear whether T cell expression of BCL6 is required to support the formation and expansion of anti-insulin B cells in the repertoire (initial B cell immune tolerance breach), anti-insulin B cell licensing by anti-insulin T cells, or both. To focus on this latter possibility (which would support BCL6 targeting in human T1D after B cell autoimmunity for insulin/islet autoantigens is established), we deploy the VH125^SD^.NOD model, in which a 1-3% population of insulin-binding B cells forms in the bone marrow that seeds the periphery and supports accelerated diabetes onset ^41^. The non-IgH locus-targeted version of this BCR transgene, VH125Tg, which is not subject to BCR somatic hypermutation, also supports accelerated diabetes onset in NOD mice ^42^. This indicates that subsequent T cell selection/affinity maturation is dispensable for anti-insulin B cell expansion and pathogenic effector function as diabetogenic APCs in the VH125 model. Furthermore, a 1-3% population of insulin-binding B cells forms in the periphery of non-autoimmune, T1D-protected VH125^SD^.C57BL/6 mice, which lack the critical IA^g7^ MHC class II molecule necessary for proinflammatory islet-reactive T cell activation and spontaneous diabetes development ^43^, further highlighting the T cell independence of insulin-binding B cell formation in this BCR transgenic model. Thus, the VH125^SD^.NOD model disconnects pro-pathogenic B cell clone formation in the repertoire from subsequent anti-insulin B cell engagement of pro-pathogenic T cells to drive islet attack.

To address whether CD4-targeted *Bcl6* elimination prevents diabetes even when such a “T1D-poised” B cell repertoire has formed, we generated VH125^SD^.Bcl6^fl/fl^.NOD (termed VH125^SD^.NOD hereafter) and VH125^SD^ Bcl6^fl/fl^.Cre*^CD4^*.NOD (termed VH125^SD^ *Bcl6*^ΔCD4^) mice. We confirmed the expected presence of a 1-3% population of anti-insulin B cells (amongst total B cells) in the spleen of VH125^SD^.NOD mice (**Figure 1A**), as previously shown ^41^. Despite the presence of anti-insulin B cell populations in peripheral organs, T cell loss of *Bcl6* in VH125^SD^ *Bcl6*^ΔCD4^ mice led to nearly complete diabetes protection relative to control VH125^SD^.NOD mice **(Figure 1B).**

**Figure 1:**
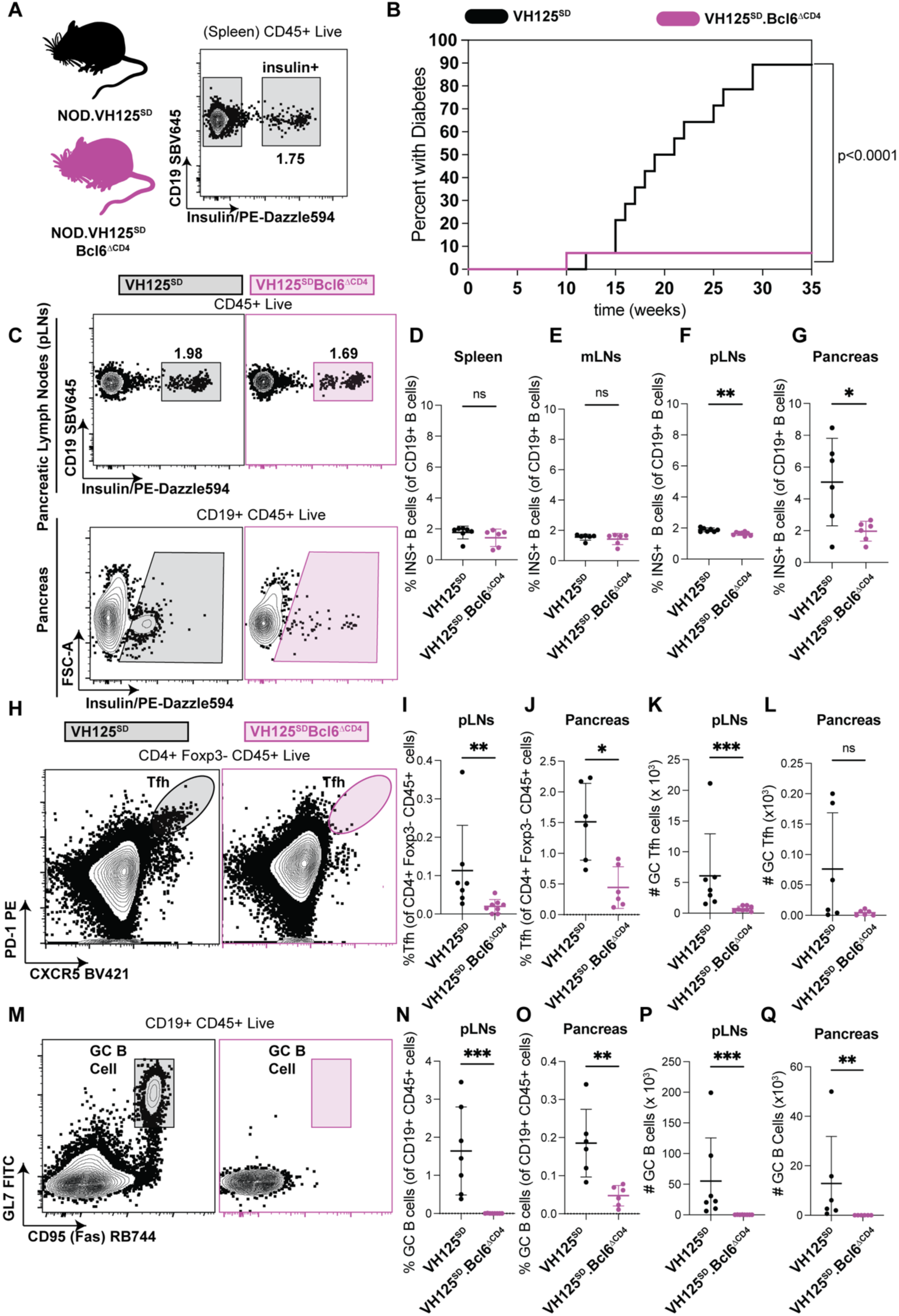
BCL6 in T cells promotes spontaneous GC formation, anti-insulin B cell infiltration of islets, and spontaneous diabetes development in VH125^SD^.NOD mice. Cells from spleen, pLNs, mLNs, and pancreas were isolated from 8-12-week-old VH125^SD^.NOD and VH125^SD^.Bcl6^ΔCD4^.NOD mice (genotypes fully defined in Methods). **(A)** A representative flow cytometry plot from spleen shows the frequency of insulin-binding B cells (Insulin+) identified using biotinylated human insulin/streptavidin fluorochrome as in Methods amongst total B cells (live singlet CD45^+^ CD19^+^ lymphocytes). **(B)** Diabetes was monitored in cohorts of female VH125^SD^.NOD mice (n = 14, black line) and VH125^SD^. Bcl6^ΔCD4^.NOD littermates (n = 14, pink line) from 10-35 weeks of age. Mice were considered diabetic after two consecutive blood glucose readings >250mg/dl, p < 0.0001, log-rank test. **(C-G)** Representative flow cytometry plots of pancreatic draining lymph nodes and pancreata gating on anti-insulin B cells as in **(A)** and as described in Methods for pancreas are shown **(C)**. The frequency of anti-insulin B cells (amongst total B cells) in **(D)** spleen, **(E)** mesenteric lymph nodes (mLNs), **(F)** pancreatic draining lymph nodes, and **(G)** pancreata are plotted for individual mice of the indicated genotypes. **(H-L)** Representative flow cytometry plots of **(H)** Tfh cells (live singlet CD45^+^ CD4^+^ PD-1^hi^ CXCR5^hi^ Foxp3^-^ lymphocytes) are shown with **(I-J)** frequencies amongst total CD4^+^ cells and **(K-L)** numbers of Tfh cells in pLNs and pancreata plotted for individual mice. **(M-Q)** Representative flow plots of **(M)** GC B cells (live singlet CD45^+^ CD19^+^ Fas^+^ GL7^+^ lymphocytes) with **(N-O)** frequencies of GC B cells amongst total B cells and **(P-Q)** numbers shown for pLNs and pancreata. **(C-Q)** n = 6-8 mice per group, n = 5 independent experiments, Mann-Whitney U test, bars representative of mean +/- standard deviation. * p < 0.05, ** p < 0.01, ***p < 0.001

We next examined insulin-binding B cell phenotypes in pancreatic lymph nodes and pancreas via flow cytometry (**Figure 1C)**. There was no observable difference in the proportion of anti-insulin B cells in spleen or mesenteric lymph nodes (**Figure 1D-1E**), with a modest reduction in pancreatic lymph nodes noted between the *Bcl6*-sufficient and deficient groups (**Figure 1F**). However, a significantly reduced frequency of insulin- binding B cells was observed in the pancreas of *Bcl6*-deficient (vs. *Bcl6*-sufficent) mice (**Figure 1G**). Total B cell and CD4+ T cell frequencies were unchanged regardless of *Bcl6* presence or absence **(Supplemental Figure 1)**. Splenic B cell developmental populations in VH125^SD^ mice were unperturbed by *Bcl6* loss in T cells, with no changes noted in transitional (T1 or T2), follicular, pre-marginal zone, or marginal zone populations (**Supplemental Figure 2)**.

The *Bcl6*-deficient group showed a reduced proportion and number of Tfh and GC B cells (gated as in **Figure 1H** and **1M**, full gating scheme in **Supplemental Figure 3**) in pancreatic lymph nodes and pancreas relative to *Bcl6*-sufficient mice (**Figure 1H-1Q**), with the exception that pancreatic Tfh cell numbers trended down but were not significantly different **(Figure 1L**). T follicular regulatory cells (Tfr), which regulate the GC response ^44^, were significantly reduced in the *Bcl6*-deficient group in spleen and pLNs, with the exception of pancreata in which Tfr are rare (**Supplemental Figure 4A- B).** CD4+ T peripheral helper (Tph) populations, defined as PD-1^hi^ ICOS^hi^, showed no significant differences with T cell loss of BCL6 in spleen and pancreatic draining lymph nodes, with a downward trend noted in pancreata **(Supplemental Figure 4C-D).** Thus, T cell loss of *Bcl6* prevents diabetes in VH125^SD^.NOD mice despite the expanded pool of anti-insulin B cells that reach the periphery.

### BCL6 loss in T cells reduces insulitis severity in VH125^SD^.NOD mice but does not impact T/B lymphocyte organization

Given the diabetes protection observed, we next examined whether T cell expression of BCL6 was required for pancreatic islets infiltration by T and B lymphocytes, which we evaluated in 8-16-week-old VH125^SD^.NOD mice, in line with insulitis kinetics in this model (**Figure 2A**) ^41^. Average insulitis was reduced in 8-12-week-old VH125^SD^ Bcl6^ΔCD4^ mice, relative to *Bcl6*-sufficient controls, with a corresponding increase in the percentage of islets with no insulitis present in the *Bcl6*-deficient group (**Figure 2B-D**).

**Figure 2:**
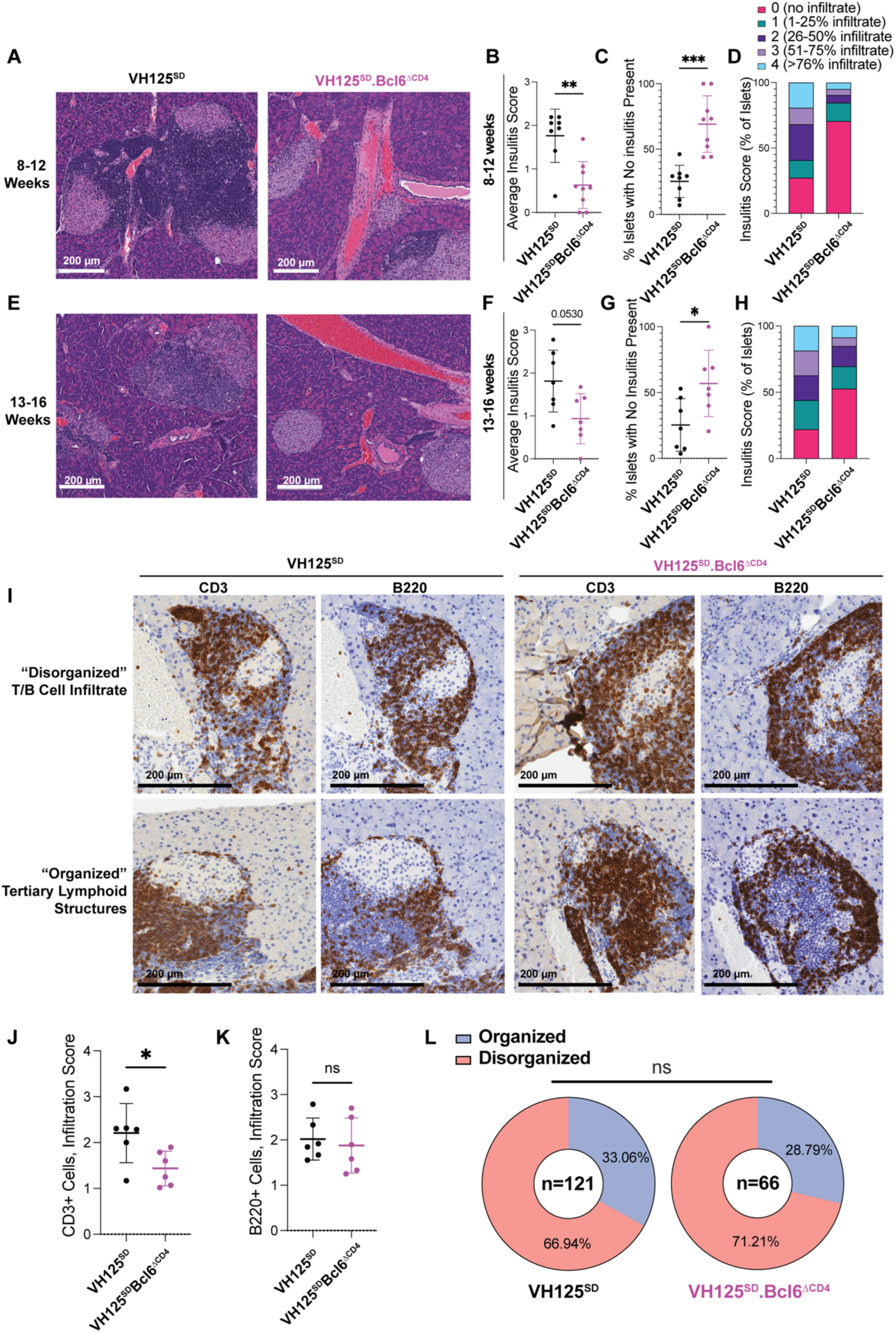
T cell expression of BCL6 supports enhanced insulitis severity, but is dispensable for tertiary lymphoid structure organization in VH125^SD^.NOD mice. Pancreata were harvested from female, pre-diabetic VH125^SD^.NOD and VH125^SD^Bcl6^ΔCD4^.NOD mice at 8-12 and 13-16 weeks of age and were formalin fixed, and paraffin embedded. **(A-H)** 10 µm pancreas sections were stained with hematoxylin and eosin (H&E) and blind scored, with individual mice plotted. **(A & E)** Representative H&E-stained sections are shown, with arrows pointing to islets. All islets were blind scored, with average insulitis scores shown for control VH125^SD^.NOD mice (black) and VH125^SD^Bcl6^ΔCD4^.NOD mice (purple) at **(B)** 8-12 weeks and **(F)** 13-16 weeks. The percentage of islets which had no lymphocytic infiltrate present were calculated for **(C)** 8-12 weeks and **(G)** 13-16 weeks. **(D and H)** The percentage of islets with each score for all pancreata for 8-12 and 13-16 weeks is shown. **(I-L)** Pancreas sections from the 8- 16-week-old cohort that had the highest insulitis infiltrate (n=6 mice per group) were obtained and stained with antibodies reactive against CD3 (T cells) or B220 (B cells). **(I)** Representative images show “disorganized” T cell-B cell infiltration (top), and “organized” TLS (bottom). Islets were scored separately for T cell (CD3) and B cell (B220) infiltrate as follows: 0 (no T/B infiltrate), 1 (>25% infiltrate), 2 (25-50% infiltrate), 3 (50-75% infiltrate), and 4 (>75% infiltrate). Average infiltration score for **(J)** CD3+ and **(K)** B220+ cells in islets is shown. **(L)** Infiltrated islets that scored 2 or above were blind scored as organized (blue) or disorganized (red) in both VH125^SD^.NOD mice and VH125^SD^Bcl6^ΔCD4^.NOD, n = number of islets scored. **(A-L)** n = 6-8 mice per group, Bars represent mean +/- standard deviation, ns = not significant, * p < 0.05, ** p < 0.01, *** p < 0.001, **(B-C, F-G, J-K)** Mann Whitney U test or **(L)** chi-square test.

The magnitude of this decrease was less apparent in 13-16-week-old mice, an interval at which 5-10% of mice typically develop diabetes **(Figure 2E-2H)**.

Organized T and B cell zones, termed tertiary lymphoid structures (TLS), can form in the pancreatic islets of NOD mice and contain GCs ^45^. To examine whether a pre-existing repertoire bias towards anti-insulin B cell formation alters TLS formation and organization, we performed immunohistochemistry to detect B and T lymphocytes in pancreas sections (as outlined in Methods and shown in **Figure 2I**). CD3 T cell infiltration was reduced with loss of *Bcl6* **(Figure 2J),** while B cell infiltration was largely similar between both genotypes **(Figure 2K)**, with some organized TLS still noted in both genotypes (**Figure 2L)**. Thus, T cell loss of BCL6 failed to eliminate TLS organization in a model where insulin autoreactive B cell expansion is present.

### T cell expression of BCL6 promotes proliferation, activation, and upregulation of costimulatory ligands on insulin-binding B cells in T1D-associated organs

Anti-insulin B and T cells can drive each other to proliferate *in vitro* and can push each other to adopt GC B cell and GC Tfh fates, respectively, *in vivo* ^11,19^. We therefore sought to address whether *Cd4*-Cre-mediated *Bcl6* loss impacted spontaneous anti-insulin B cell proliferation *in vivo*. Given that GC B cells are highly proliferative ^46^, we examined the frequency of Ki67+ cells amongst non-GC B cells to eliminate this potential bias, as in **Figure 3A-B**. While few non-insulin-binding B cells were Ki67+, insulin-binding B cell proliferation (marked by Ki67 positivity) was elevated by comparison, which was reduced in the *Bcl6*-deficient group in pancreatic lymph nodes (**Figure 3C**) and pancreas (**Figure 3D**).

**Figure 3:**
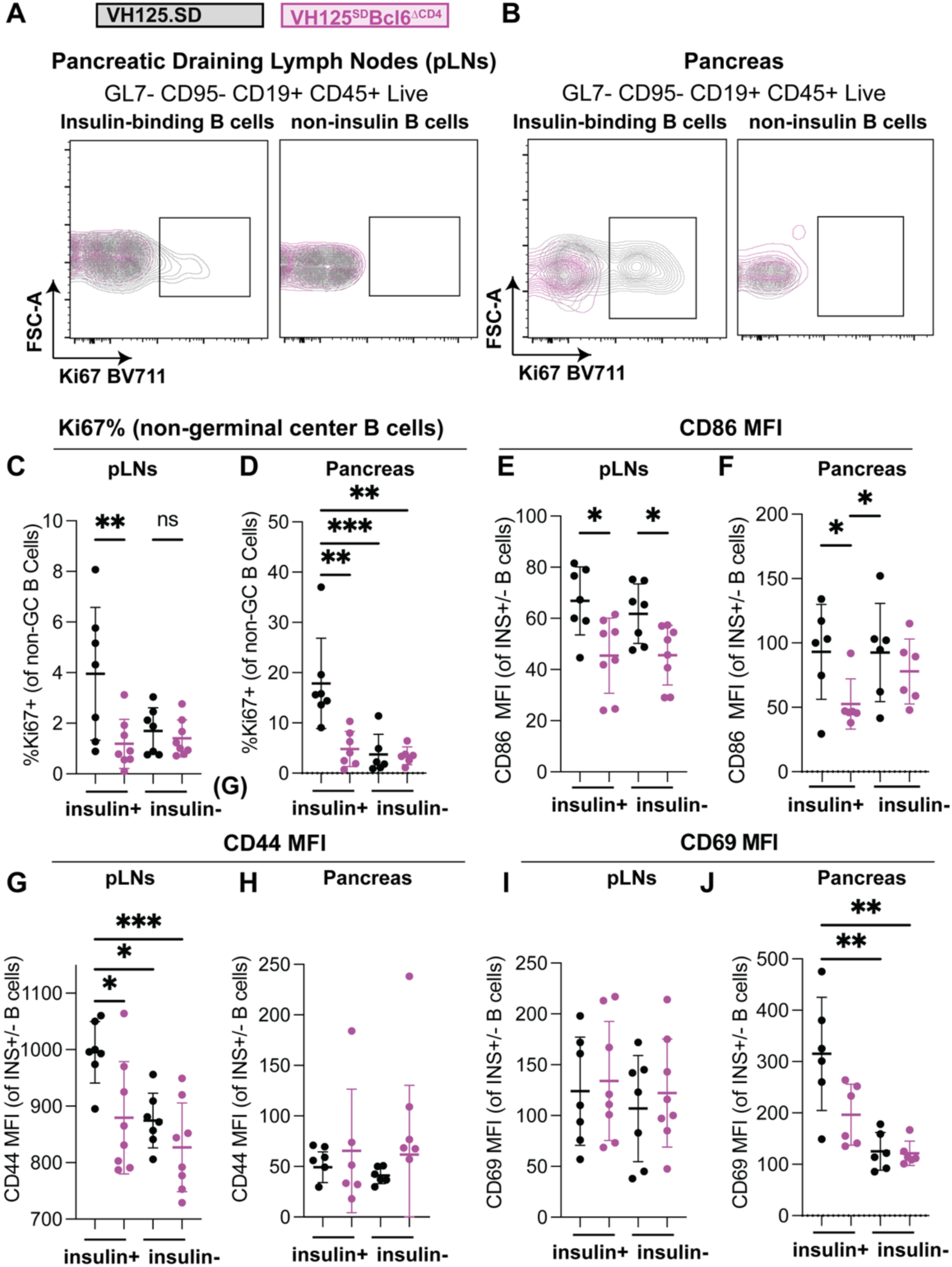
BCL6 in T cells increases activation and proliferation markers of insulin-binding B cells relative to non-insulin-binding B cells. Cells were isolated from 8-12-week-old, female, pre-diabetic VH125^SD^.NOD with and without *Cd4*-Cre *Bcl6* deletion from **(A-D)** pancreatic lymph nodes (pLNs) and **(E-H)** pancreata. **(A, E)** Representative flow cytometry plots show Ki67 staining overlays of insulin-binding (left) or non-insulin-binding (right) B cells (identified as in Figure 1) from each genotype within live singlet CD45^+^ CD19^+^ lymphocytes. **(B-D)** Non-GC (Fas^-^ GL7^-^) B cells were further gated on insulin-binding (ins+) and non-insulin-binding (ins-) and the frequency of cells that were **(A, E)** Ki67+ (a marker of proliferation), **(B, F)** CD86 (T cell co-stimulatory molecule), **(C, G)** CD44 (activation marker), and **(D, H)** CD69 (activation marker) in the pLNs (top row) and pancreas (bottom row). **(C-H)** n = 6-8 mice per group, Kruskal-Wallis test with post-hoc uncorrected Dunn’s test of multiple comparisons. * p < 0.05, ** p < 0.01, *** p < 0.001. Bars represent mean +/- standard deviation.

Insulin-binding B cells upregulate the T cell costimulatory molecule, CD86, relative to non-insulin-binding B cells in the pancreas, but not in the spleen of VH125Tg.NOD mice^30^. This is presumably related to their pathogenic function as antigen-presenting cells in T1D ^11,41^. We therefore assessed markers of co-stimulation (CD86) and activation (CD44 and CD69) based on insulin specificity and *Bcl6* genotype. CD86 levels were reduced in insulin-binding B cells in VH125^SD^Bcl6^ΔCD4^ mice compared to control VH125^SD^ mice in both pancreatic lymph nodes (**Figure 3E**) and pancreas (**Figure 3F).** CD86 levels were also decreased in VH125^SD^Bcl6^ΔCD4^ non-insulin-binding B cell compartments in the pancreatic lymph nodes but not the pancreas (**Figure 3E-3F)**. CD44, a marker for activation, was reduced with the loss of *Bcl6* in insulin-binding B cells in pancreatic lymph nodes (**Figure 3G**) but not pancreas (**Figure 3H**). Insulin- binding B cells expressed higher levels of CD69 relative to non-insulin-binding B cells in the pancreas, which were not altered by T cell loss of BCL6 in either pancreas or pancreatic lymph nodes (**Figure 3I-J).** In contrast, few BCL6-dependent differences were seen in Ki67, CD86, CD44, or CD69 across non-insulin-binding B cell populations in either pancreatic lymph nodes or pancreas, except for CD86 levels in the pancreatic lymph nodes (**Figure 3C-3J**). Together, these results show that T cell expression of BCL6 supports anti-insulin B cell activation, expression of T cell co-stimulatory markers, and proliferation, all of which hold potential to impact their function as pathogenic antigen-presenting cells.

### IgG1 and IgG2b class switching is reduced amongst insulin-binding and non-insulin-binding B cells by T cell loss of BCL6

Although class switching occurs largely before the GC B cell identify forms, BCL6 stabilizes B cell class switching to IgG1 ^47–49^. The two major isotypes of spontaneous insulin autoantibody serologically detected in wildtype NOD mice are IgG1 and IgG2b, with predominantly IgG1 isotype ^50^. T cell loss of *Bcl6* loss in VH125^SD^.NOD mice led to a reduced frequency of IgG1+ and IgG2b+ B cells amongst insulin-binding as well as non-insulin-binding B cells in pancreatic lymph nodes **(Figure 4A-D).** Although class-switched B cells are readily detected in VH125^SD^.NOD mice, the presence of circulating IgG anti-insulin Ab is almost completely suppressed ^41,51^. Consistent with this, we detected little insulin-specific IgG autoantibody regardless of *Bcl6* genotype, but did observe a significant, albeit small reduction in anti-insulin IgG (**Figure 4E**). Thus, BCL6 loss leads to reduced spontaneous anti-insulin B cell and non-insulin-binding B cell class-switching to IgG1 and IgG2b.

**Figure 4:**
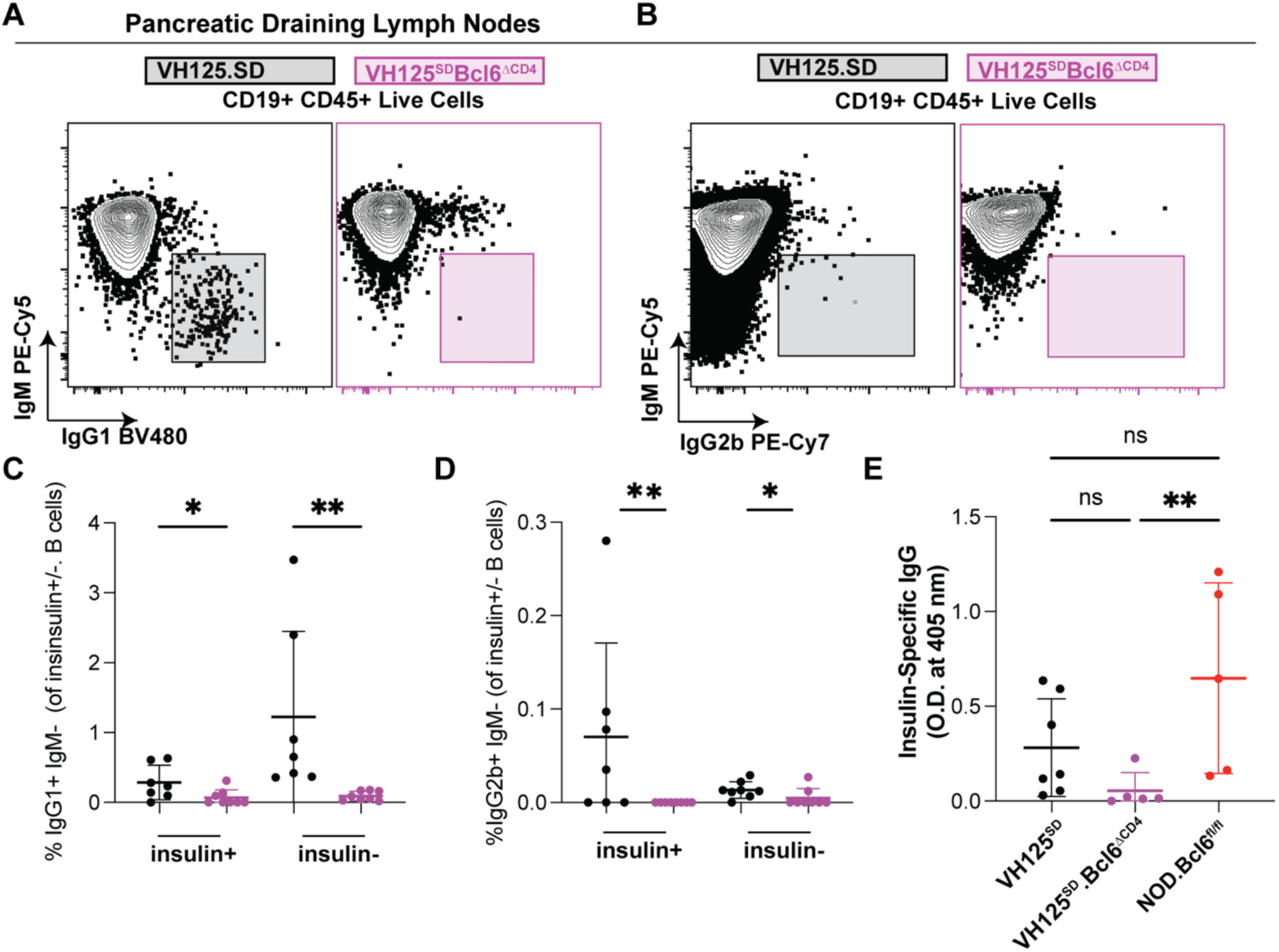
BCL6 loss in T cells prevents class switching of autoreactive B lymphocytes. Pancreatic draining lymph nodes (pLNs) were harvested from pre-diabetic VH125^SD^ mice with or without *Cd4*-Cre loss of *Bcl6*. **(A)** Class switched IgG1+ B cells were defined as IgG1+ IgM-CD19+ live cells in either insulin- or insulin+ autoreactive B cells. **(B)** Class switched IgG2b+ B cells were defined as IgG2b+ IgM-CD19+ live cells in either insulin- or insulin+ autoreactive B cells. **(C-D)** Quantification of non-autoreactive, insulin-B cells and autoreactive, insulin+ B cells in both **(C)** IgG1+ and **(D)** IgG2b+ for both VH125^SD^ (black) and VH125^SD^Bcl6^ΔCD4^ (purple) mice. **(E)** Anti-insulin IgG autoantibody levels were determined in both VH125^SD^ (black), VH125^SD^Bcl6^ΔCD4^ (purple), and non-transgenic NOD.Bcl6^fl/fl^ (red) mice via ELISA. O.D. values represent the difference in O.D. values of non-inhibited to inhibited insulin-binding. Each dot is the average value of 3 technical replicates per mouse. **(C-E)** n = 5-8 mice per group, Kruskal-Wallis non-parametric statistical tests were used with post-hoc uncorrected Dunn’s test of multiple comparisons. * p < 0.05, **p < 0.01, ns=not significant. Bars indicate mean +/- SD.

### BCL6 in T cells promotes anti-insulin B cells skewing towards atypical B cell phenotypes preferentially in T1D-associated organs

Increasing evidence points to the likely importance or even centrality of atypical B cell populations in chronic infections and auto-Ab-mediated diseases such as SLE ^32,40^. However, B cell proclivity towards atypical B cell populations has not yet been examined in the NOD mouse strain. We identified proliferative (Ki67+) anti-insulin B cells in pancreas and pancreatic lymph nodes, even when GC B cells are excluded from analysis (**Figure 3A-B**). This led us to hypothesize that anti-insulin B cells could be engaging T cells outside of GCs. We therefore examined atypical B cell markers (CD11b, CD11c, and T-bet) to determine if insulin-binding B cells skew towards atypical B cell phenotypes, with representative gating shown in (**Figure 5A, 5C**) ^36^. Insulin- binding B cells skewed towards CD11b+ CD11c+ (**Figure 5B**) and T-bet+ CD11c+ (**Figure 5D**) B cell populations in the pancreas and pancreatic lymph nodes, compared to non-insulin binding B cells (**Figure 5E)**. In contrast, we observed no differences in atypical B cell populations between insulin-binding and non-insulin-binding B cells in the spleen **(Figure 5F).** We observed *Cd4*-Cre *Bcl6*-dependent loss of atypical B cell populations in both pancreatic draining lymph nodes and pancreas, but not spleen (**Figure 5A-F)**. Overall, these data show that insulin-binding B cells skew towards CD11b+ CD11c+ and Tbet+ CD11c+ atypical B cell subsets in T1D-associated organs, with some subsets (CD11b+ CD11c+) reduced by T cell loss of *Bcl6*.

**Figure 5:**
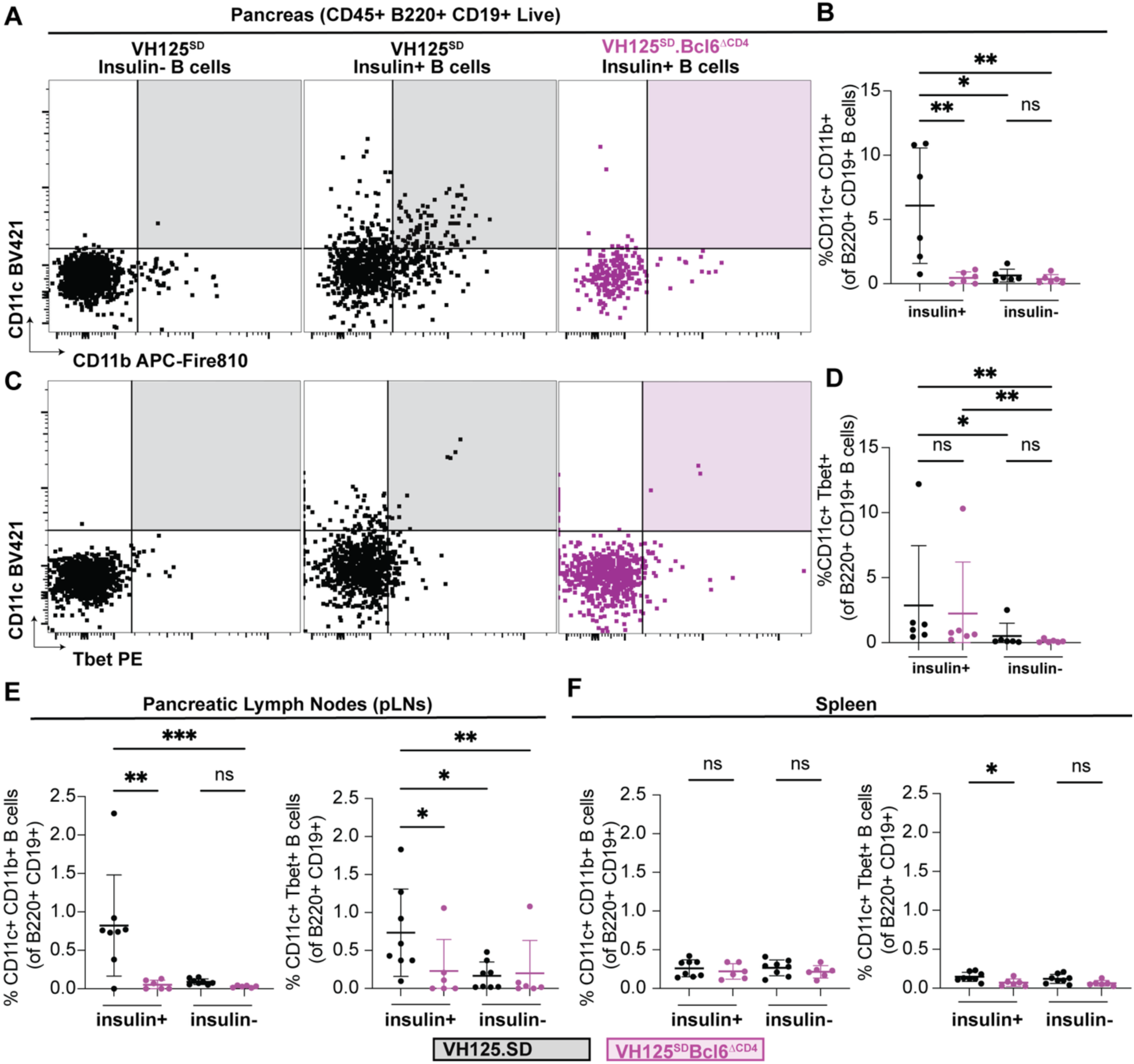
Insulin-binding B cells skew towards CD11c^+^ T-bet^+^ and CD11b^+^ CD11c^+^ atypical B cell subsets, some of which are reduced by loss of *Bcl6*. Spleen, pancreatic draining lymph nodes, and pancreata were harvested from pre-diabetic, 8- 12-week-old VH125^SD^.NOD mice. **(A)** Representative flow plots show CD11c+/T-bet+ cells amongst B220+ CD19+ live singlet lymphocytes for both non-insulin autoreactive (insulin-, left) and insulin autoreactive (insulin+, middle, right) B cell populations in the pancreas. **(B)** Quantification of the proportion of CD11c+ T-bet+ B cells in pancreata for both insulin- and insulin+ B cells for VH125^SD^ (black) and VH125^SD^Bcl6^ΔCD4^ (purple) mice. **(C)** Representative flow plots of CD11c+/CD11b+ cells amongst B220+ CD19+ live singlet lymphocytes for both insulin- and insulin+ B cell populations in the pancreas. **(D)** Quantification of the proportion of CD11c+/Tbet+ cells amongst both insulin- and insulin+ B cells in the pancreas of VH125^SD^ (black) and VH125^SD^Bcl6^ΔCD4^ (purple) mice. **(E-F)** Quantification of the proportion of CD11b+/CD11c+ and CD11c+/Tbet+ B cells in **(E)** pancreatic draining lymph nodes and **(F)** spleen for both insulin- and insulin+ B cells for VH125^SD^ (black) and VH125^SD^Bcl6^ΔCD4^ (purple) mice. **(A-F)** Kruskal Wallis test with post-hoc uncorrected Dunn’s multiple comparison test, n = 6-8 mice per group. * p < 0.05, ** p < 0.01, *** p < 0.001, ns = not significant.

### B cell populations expressing activated, GC, atypical, and atypical memory phenotypic markers depend on T cell expression of BCL6

To avoid user bias in identifying B cell changes when BCL6 is lost by T cells, we next used the minimally supervised clustering tool, Tracking-Responders Expanding (T-REX). For example, T-REX combined with Marker Enrichment Modeling (MEM) identified a human T cell population that expanded in a SARS-CoV-2 vaccine response which was highly enriched for antigen-reactive T cells ^52–54^. T-REX identified seven populations of cells in VH125^SD^.NOD pancreata that contracted (blue/light blue) with T cell loss of BCL6 (**Figure 6A**). MEM identified GC markers GL7, Ki67, and CD95 in Cluster 5, markers expressed by memory B cells, such as CD80, CD73, and PDL2 in Cluster 2, 3, and 4 (with activation and atypical B cell markers in Cluster 4), and an anti-insulin population in Cluster 1 and 7 (**Figure 6B)**. Heatmap examination of Clusters 1 and 7 (anti-insulin B cell populations) revealed increases in some markers of atypical and memory B cell markers (including CD11c, CD11b, PDL2, and CD80) not included in the initial MEM labelling (**Figure 6C-D)**. Heatmap examination of Cluster 3 further revealed increases in some markers of atypical memory B cell markers (including CD11c, CD11b, PDL2, and CD80) (**Figure 6D)**. Similar populations were contracted in pancreatic draining lymph nodes by T cell loss of BCL6 (**Supplemental Figure 5-A-C**). Thus, T-REX confirmed changes in some populations identified through conventional gating analysis (e.g., GC B cells, Cluster 1) and identified another population (CD80hi PDL2hi, Cluster 2) that was not initially evaluated.

**Figure 6.**
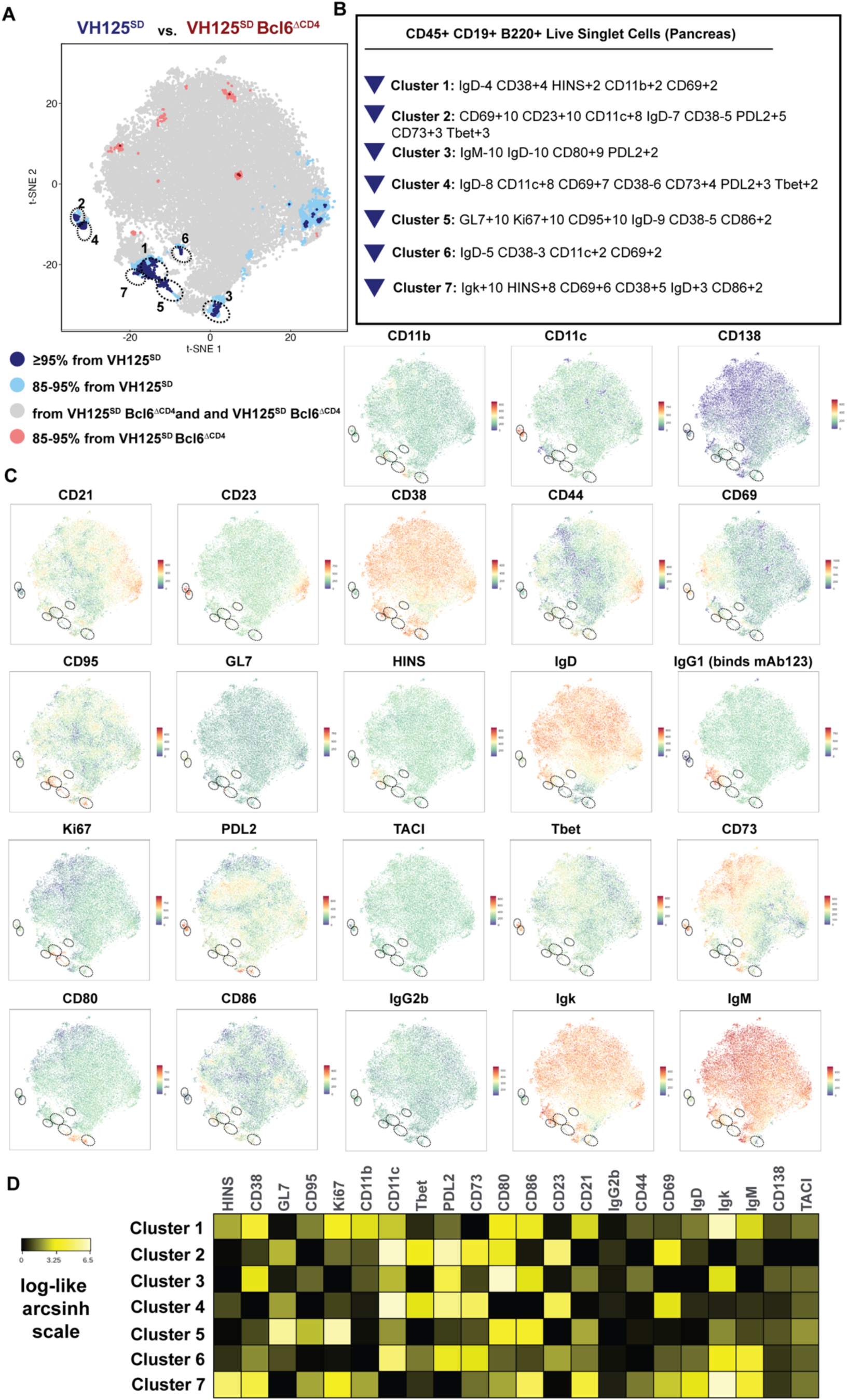
BCL6 loss in T cells leads to contracted pancreatic B cell populations with phenotypic attributes of GC, activated, and atypical memory B cells in VH125^SD^.NOD mice. Pancreata were harvested from pre-diabetic, 8-12-week-old VH125^SD^.NOD mice. B220+ CD19+ CD45+ live singlets from n = 12 mice were normalized via CyCombine and concatenated into two groups, *Bcl6*-sufficient (VH125^SD^) and *Bcl6*-deficient (VH125^SD^Bcl6^ΔCD4^.NOD). **(A)** t-SNE was used to perform dimensionality reduction based on phenotypic marker expression profiles. Following this, the minimally supervised analysis tool, Tracking Responders Expanding (T-REX) was used to identify populations that were increased (red: > 95% or peach: 85-95% population change) or decreased (blue: > 95% or light blue: 85-95% population change) by *Cd4*-Cre loss of *Bcl6* in this model, as illustrated in the t-SNE plot. **(B)** Marker Enrichment Modeling (MEM) labels show the phenotypic marker features that define each of the contracted populations. Features enriched by at least +2 or -2 on a scale from 0 to 10 are shown. **(C)** A rainbow intensity scale indicates expression levels of the phenotypic markers indicated at the top of each t-SNE plot with red representing high levels and blue representing low levels of expression. CD45, B220, CD19, and the viability dye expression was omitted from the t-SNE analysis given these markers were used to gate on the parent population that went into phenotypic clustering via t-SNE and T-REX visualization and analysis. Note that mAb123, which binds to insulin-binding B cells, also binds to IgG1 antibody, thus providing better separation of insulin-binding B cell populations. **(D)** Heatmap shows relative expression of the indicated markers for each cluster defined as in *(B)* using a log-like arcsinh scale and appropriate arcsinh factors.

### T cell expression of BCL6 supports insulin-binding B cell skewing towards an atypical memory-like population

We next performed minimally supervised analysis using T-REX after gating on insulin- binding B cells to potentially uncover additional phenotypic changes caused by CD4- driven *Bcl6* loss; however, no significantly expanded or contracted clusters were identified (**Supplemental Figure 6**). Given that we observed phenotypic changes in anti-insulin B cells with classical flow cytometry gating (**Figures 3-5)**, we also performed t-SNE-based clustering, which defined four phenotypic clusters in the pancreas (**Figure 7A**). Cluster 1 was significantly reduced in VH125^SD^Bcl6^ΔCD4^ mice relative to VH125^SD^ controls, while Clusters 3 and 4 trended upwards (**Figure 7B**). Cluster 1 exhibited increased Ki67, PDL2, CD80, and T-bet, suggesting this may be a proliferative, atypical memory B cell cluster (**Figure 7C-D**). PDL2 and CD80 are not exclusively found on memory B cells, thus future studies are required to determine whether this population is memory. Nonetheless, in sharp contrast to cluster 1, clusters 2-5 were unaffected by BCL6 loss in T cells (**Figure 7B**). Pancreatic draining lymph nodes exhibited a reduction of an atypical-memory-like B cell population with T cell loss of BCL6 (**Supplemental Figure 7A-D)**. Overall, this study identified several ways in which BCL6-expressing T cells redirected insulin-binding B cells towards pro-pathogenic fates, which were associated with diabetes development.

**Figure 7:**
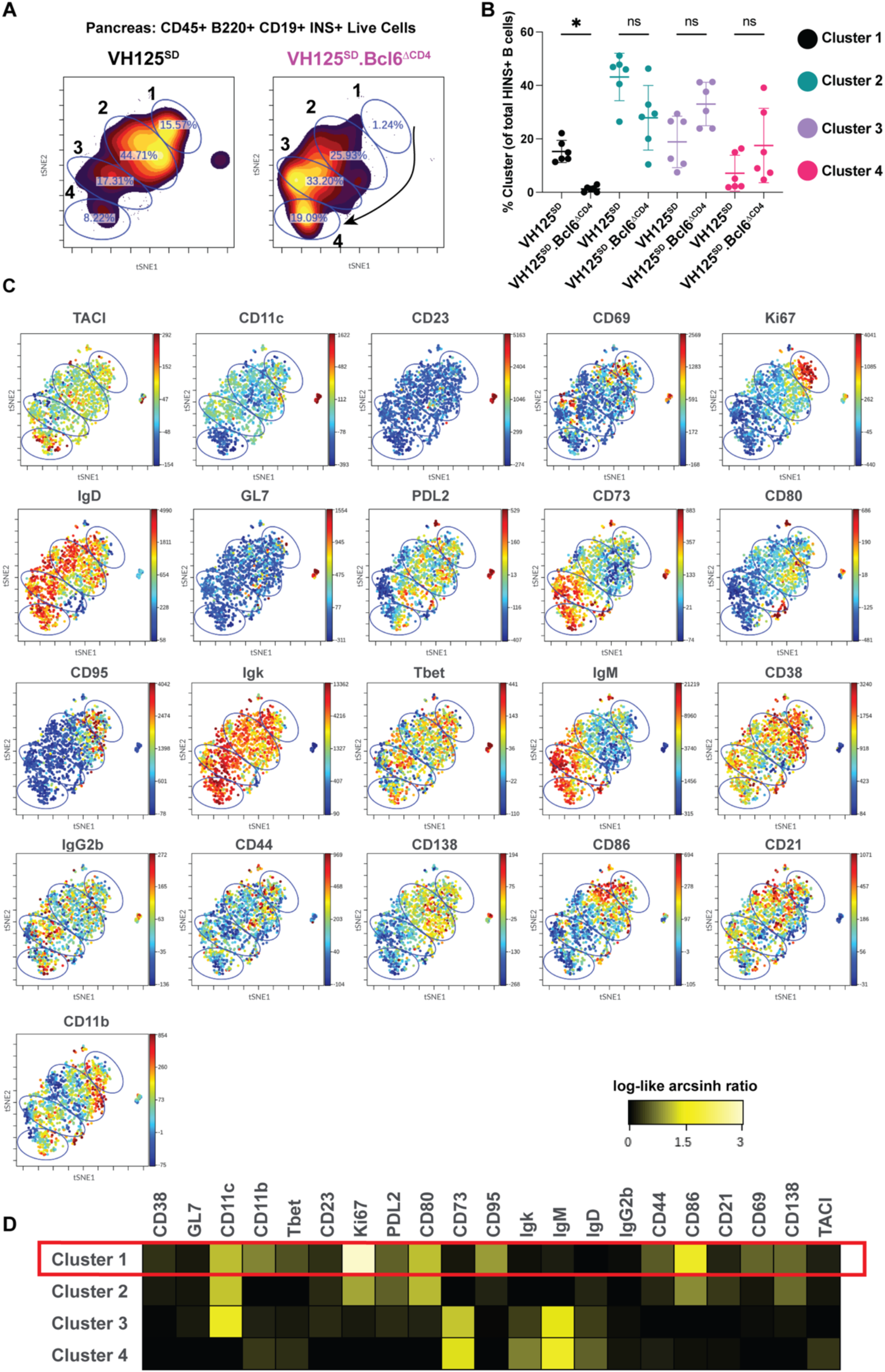
BCL6 loss in T cells promotes formation of a phenotypically defined atypical memory B cell population in the pancreas of VH125^SD^.NOD mice. Pancreata were harvested from pre-diabetic, 8–12-week-old VH125^SD^.NOD mice. Insulin-binding B220+ CD19+ CD45+ live singlets cells (n = 6-7 mice per genotype) were normalized via CyCombine and concatenated into two groups, *Bcl6*-sufficient (VH125^SD^) and *Bcl6*-deficient (VH125^SD^Bcl6^ΔCD4^.NOD). **(A)** t-SNE was used to perform dimensionality reduction based on phenotypic marker expression profiles. Clusters were manually defined as indicated. **(B)** The cluster frequency is shown for each genotype, with individual mice plotted and means indicated. * p < 0.05, Kruskal-Wallis test with post-hoc uncorrected Dunn’s test. **(C)** A rainbow intensity scale indicates expression levels of the phenotypic markers indicated at the top of each t-SNE plot. Insulin-binding, CD45, B220, CD19, and viability dye markers were omitted from the t-SNE analysis given they were used in parent population gating upstream of t-SNE analysis. **(D)** Heatmap shows relative expression of the indicated markers for each cluster defined as in *(A)* using a log-like arcsinh scale and appropriate arcsinh factors.

## Discussion

In this study, we find that T cell loss of BCL6 impacts the fate of anti-insulin B lymphocytes, limiting their upregulation of CD86, proliferation, IgG switching, GC B cell differentiation, and skewing towards atypical B cell populations. These changes culminated in near complete diabetes prevention in VH125^SD^.NOD mice, which otherwise develop accelerated diabetes when T cells are competent to express BCL6. Our findings suggest a direct role of BCL6-expressing CD4 (and potentially CD8) T cells in licensing pathogenic B cell subsets, which in turn support T cell-mediated beta cell attack. Clinical immunotherapy efforts increasingly focus on intervening during the pre- diabetic interval ^3^. Stage 1 T1D is typically defined by pre-existing B lymphocyte autoimmunity against islet autoantigens (as indicated by islet autoantibody seropositivity). These findings highlight the translational potential of targeting BCL6 in T1D, as BCL6 loss in T cells provides robust protection against T1D despite the forced generation of insulin-binding B cells in this BCR transgenic model. Furthermore, BCL6 inhibitors are currently in clinical trials for treatment of B-cell Non-Hodgkin’s Lymphoma, streamlining the process for regulatory approval as a new potential T1D immunotherapy^55^. Our findings clarify that *Cd4*-Cre-mediated *Bcl6* loss not only abrogates GC B cell and Tfh formation but also reduces the proliferative and T cell costimulatory potential of anti-insulin B cells, suggesting a direct role of BCL6-expressing CD4 (and potentially CD8) T cells in licensing pathogenic B cell subsets, which in turn support T cell-mediated beta cell attack.

We observed an increased frequency of anti-insulin B cells in the pancreas (relative to spleen or pancreatic lymph nodes) which was reduced by T cell loss of *Bcl6*. We found that anti-insulin B cells selectively upregulated CD69 ∼two-fold in the pancreas of VH125^SD^.NOD mice, with an increased skewing towards a proliferative phenotype (Ki67) relative to non-insulin-binding B cells, which was ablated by T cell loss of BCL6. These data are consistent with anti-insulin B cells receiving reduced T cell help with loss of BCL6. Our findings also demonstrate reduced IgG1 and IgG2b class switching in VH125^SD^.Bcl6^DCD4^ vs. VH125^SD^ mice, which is consistent with an established role for BCL6 in promoting class-switch recombination, particularly to IgG subclasses ^48^.

Amongst islets containing moderate to heavy T and B cell infiltrates in NOD mice, ∼50% of have organized structures with discrete T and B cell zones ^21,27^. Of note, the requirement for TLS in T1D development is challenged by experiments showing that anti-CXCL13 treatment blocks organized TLS and GC formation in islets but fails to prevent diabetes in this model ^56^. Here, we show that organized T/B cell infiltrates were less prevalent in the VH125^SD^.NOD model, with only ∼30% being classified as organized by the same criteria. TLS were still observed despite T cell loss of BCL6 in VH125^SD^.NOD mice, suggesting that T cell expression of BCL6 is not required for these organized T/B zones to form. The reduced TLS organization observed may reflect the increased frequency of insulin-binding B cells available outside of TLS/GCs to provide T cell help, which will require future studies to evaluate.

Atypical B cells have been implicated in other autoimmune diseases ^32,57^, yet this is the first study to highlight their increased presence in the NOD model of T1D. We observed significant enrichment of insulin-binding B cells amongst T-bet^+^ CD11c^+^ and CD11b^+^ CD11c^+^ populations in the pancreas and pancreatic lymph nodes of T1D-prone VH125^SD^.NOD mice, with T cell loss of *Bcl6* chiefly impacting skewing to the CD11b^+^ CD11c^+^ subset. Future experiments are needed to highlight the potential role, migration, and differentiation, of these subsets.

Limitations of our study include the following. Our *Cd4*-Cre system removes *Bcl6* in both CD4 and CD8 T cells ^58^. Additional experiments are therefore required to determine the individual contributions of BCL6 in specific CD4, CD8, and other immune populations in the context of T1D pathogenesis. Insulin-binding B cell upregulation of T cell co-stimulatory molecules (and potentially their potency as antigen-presenting cells) was impacted by T cell expression of BCL6. However, we did not evaluate the functional impact of this change via B cell antigen presenting cell assays, or how the generation of proinflammatory, antigen-specific T cells might be modified by T cell loss of *Bcl6*.

One currently unexplored possibility is whether the molecular regulation of atypical B cells is altered in infection versus autoimmunity. T cell loss of *Bcl6* blocks both GC and CD11c+ T-bet+ B cells, thus it’s possible the disease protection observed in VH125^SD^.Bcl6^DCD4^ mice is related to crippled extrafollicular responses related to driving proinflammatory T cell differentiation and beta cell attack. Our work does not show where these atypical B cells originate from or how T cell loss of *Bcl6* might impact their migration or development. Future work is therefore needed to fully explore these remaining questions. Additionally, while populations of insulin-binding B cells were identified that expressed PDL2, CD80, and/or CD73 markers that are associated with memory B cell populations^59^ (sometimes in combination with atypical B cell markers), additional evidence will be required to confidently call any of the phenotypically defined populations outlined here as memory and atypical memory B cells.

T-REX was designed for analysis of paired samples ^52^, which, while successfully applied here to identify population differences amongst total B cells in non-paired samples, did not unearth all phenotypic population differences amongst insulin-binding B cells that were uncovered through “expert gating” analysis of flow cytometry data. We attribute this lack of cluster detection to insufficient statistical power to detect differences given the limited number of insulin-binding B cells that are present in these organs. Nonetheless, this work highlights the important translational potential and role of GCs and atypical B cells in T1D.

Overall, our study identifies BCL6 as a pivotal transcriptional regulator, which, when lost in T cells, limits anti-insulin B cell activation, proliferation, IgG class switching, GC B cell differentiation, and skewing into the CD11b^+^ CD11c^+^ atypical B cell subset in T1D. Given the robust protection against diabetes development observed with T cell loss of BCL6 despite persistence of insulin-binding B cells in the periphery, our findings highlight the translational potential of BCL6 inhibition in T1D, even at a point when B cell autoimmunity for insulin has already manifested, such as in Stage 1 T1D, which will require clinical trial confirmation.

## Supporting information

Supplemental Data Sheet

## Acknowledgements

This work was supported by Breakthrough T1D (formerly the Juvenile Diabetes Research Foundation) grants 2-SRA-2023-1453-S-B (RHB) and 3-PDF-2024-1495-A-N (DHM), and NIH grants F30 DK145130 (LMC), F31 DK141224 (LEB), T32 GM007347 (LMC, CMN), T32 AR059039 (TWJ), R01DK105550 (JCR). We also acknowledge the Vanderbilt University Medical Center (VUMC) Flow Cytometry Shared Resource [supported by supported by the Vanderbilt Ingram Cancer Center (P30 CA068485) and the Vanderbilt Digestive Disease Research Center (P30 DK058404)], the VUMC Tissue Pathology Shared Resource [supported by NCI/NIH Cancer Center Support Grant, P30 CA068485], and the Islet and Pancreas Analysis Core [supported by the Vanderbilt Diabetes Research and Training Center (NIH grant P30 DK20593)]. We would like to acknowledge Dr. Alexander Dent (Indiana University) for providing the original *Bcl6fl/fl*.C57BL/6/129Sv mice, which we backcrossed to the NOD strain and introgressed the VH125^SD^ transgene for these studies. We also acknowledge Dr. Mark Boothby (Vanderbilt University Medical Center) for critical manuscript review.

## Author contributions

Conceptualization, LMC, DHM and RHB; methodology, LMC, JCM, DHM, MLP, TWJ, CMN; investigation, LMC; writing – original draft, LMC, JCM, DHM, RHM; writing-review and editing, LMC, JCM, DHM, JCR, RHB; funding acquisition, LMC, DHM, LEB, JCR, RHB; resources, JCR and RHB.

## Data availability

Data acquired specifically for this study is available within the article itself and the supplemental information.

## Declaration of interests

The authors declare no competing interests. RHB has unrelated funding from the National Institutes of Health, Breakthrough T1D, argenx, and the Leona M. and Harry B. Helmsley Charitable Trust. J.C.R. is Founder and Scientific Advisor of Sitryx Therapeutics.

## Methods

### Sex as a Biological Variable

To minimize the number of mice required to power the study, all studies used female NOD mice, in line increased diabetes incidence in females relative to males in NOD mice ^60^.

### Animals

*Bcl6^fl/fl^.C57BL/6/129Sv* mice ^28^ were backcrossed at least 14 generations to the NOD background and intercrossed with *Cd4*-Cre.NOD mice (strain number: 013234; The Jackson Laboratory, Bar Harbor, ME^61^) to generate *Bcl6^fl/fl^.Cd4*-Cre^+/-^.NOD mice, as described in ^27^. This loss generates a non-functional BCL6 protein whereby the DNA-binding domain is lost, but the rest of the protein is still expressed. Bcl6^fl/fl^ *Cd4*-Cre.NOD mice were then crossed with VH125^SD^.NOD mice ^41^, in which an anti-insulin IgH transgene is targeted to the IgH locus (site-directed, SD), to generate VH125^SD+/-^.Bcl6^fl/fl^.*Cd4*-Cre^+/-^ (termed VH125^SD^.Bcl6^DCD4^) and control VH125^SD+/-^.Bcl6^fl/fl^ (termed VH125^SD^) NOD lines. All studies compared littermates to limit microbiome-related differences across groups. All mice were housed under specific pathogen-free conditions and given autoclaved food and water. These animal studies were fully approved by the AALAC-accredited Vanderbilt Institutional Animal Care and Use Committee (IACUC).

### Diabetes Monitoring

Blood glucose was measured weekly from 10-35 weeks of age in female littermates by nicking tails. Diabetes was diagnosed after two consecutive blood glucose readings >250 mg/dL.

### Histological Assessment of Insulitis and Tertiary Lymphoid Structure Organization

Mice were sacrificed and pancreata were dissected from non-diabetic female mice. Pancreata were incubated overnight in 10% formalin at room temperature and then dehydrated briefly in 70% ethanol and paraffin embedded. 10 mm sections were cut, deparaffinized, and subsequently stained with H&E by the Vanderbilt Tissue Pathology Shared Resource. Slide images were obtained using a bright field Aperio ScanScope CS. Visualization and analysis was performed by using ImageScope software (Leica Biosystems). Islets were individually blind scored for insulitis (lymphocyte invasion) as described in ^21^.

Pancreas blocks which exhibited insulitis via H&E staining above were chosen from n = 6 mice per genotype for subsequent CD3 and B220 immunohistochemical staining using a Leica Bond-Max IHC stainer. Following heat-induced antigen retrieval with Bond Epitope Retrieval Solution 2 (Leica Biosystems), 10 mm pancreas sections were stained with antibodies reactive against either CD45R/B220 (RA3–6B2) or CD3 (M-20). Islets were scored on their average infiltration of CD3 (T cells) and B220 (B cells) as described ^27^. Islets with an insulitis severity score of 2 or higher were further blind scored as “organized” (defined by organized B/T cell zones) or “disorganized” (defined by B/T infiltration that was not organized into discrete tertiary lymphoid structure zones).

### Cell Isolation from Tissues

Cells were isolated from spleen, pancreatic lymph nodes (n = 2 nodes per mouse), and mesenteric lymph nodes (n > 5 nodes per mouse) as previously described ^27^. Pancreata were promptly dissected and digested with 1 mg/mL of collagenase P (Sigma) in HBSS and incubated while shaking at 37°C for 30 minutes. The suspension was then disrupted with an 18G needle and resuspended in HBSS + 10% BCS to inactivate collagenase and cells were pelleted and resuspended in flow cytometry staining buffer (detailed below). Cell counts were performed using a Bio-Rad Tc20 Automated Cell counter.

### Antibodies and Spectral Flow Cytometry

Cells were stained for flow cytometry utilizing the murine reactive antibodies listed in **Table 1**. We previously observed that endogenous (murine) insulin “masks” insulin- binding B cell detection with biotinylated human insulin in the pancreas, but not lymph nodes or spleen ^62^. This technical issue is overcome in VH125-derived B cells by staining cells with a second antibody (mAb123) that binds a different epitope than mAb125 ^62^. Thus, for pancreas staining, cells were first incubated with human insulin (Sigma #I2643) for 30 minutes in flow cytometry (FACS) staining buffer (1X PBS + 5% BCS + 0.1% azide + 0.02% EDTA), followed by detection with mAb123-biotin/SA- fluorochrome, as described ^62^.

**Table 1.**
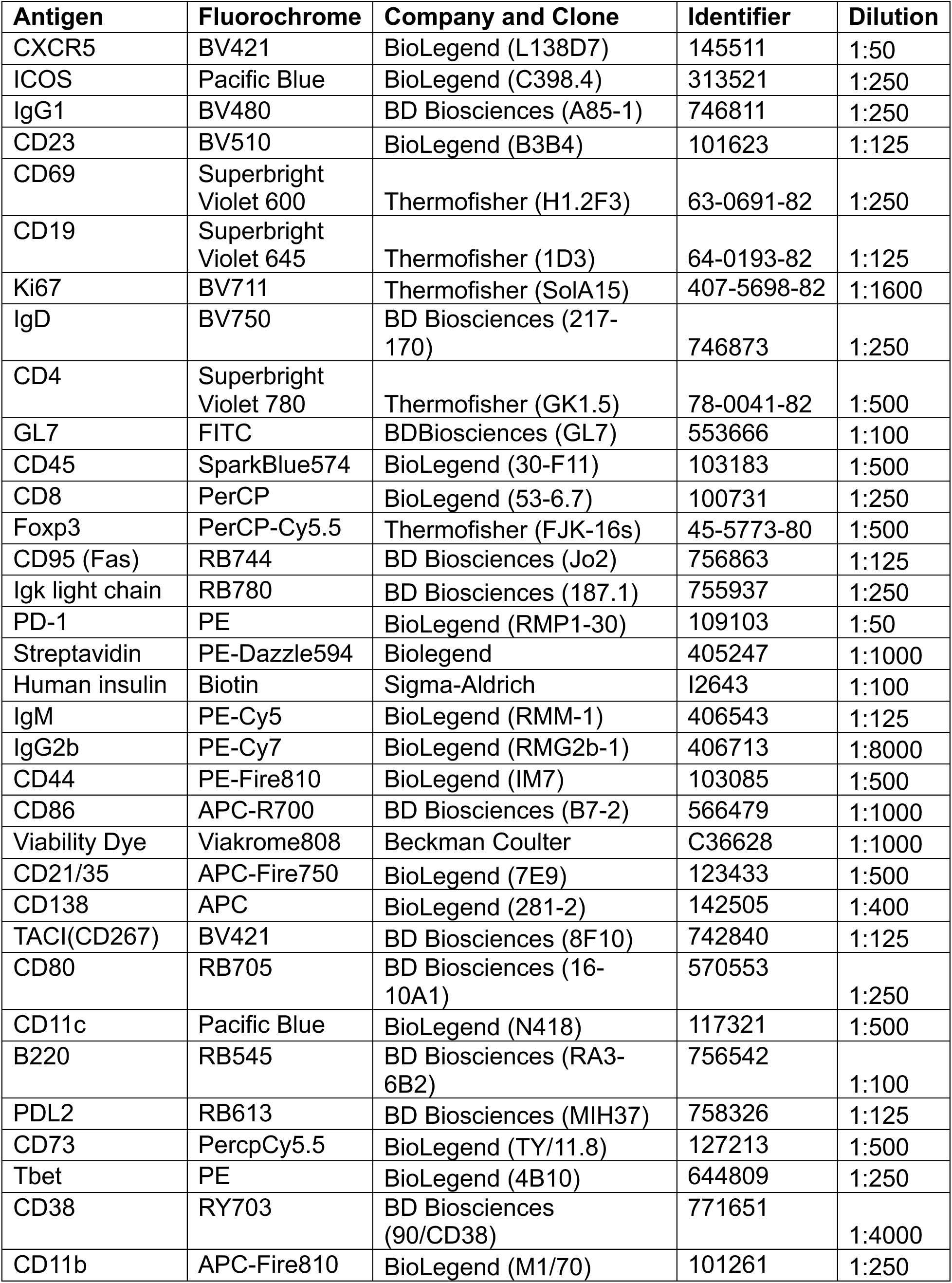
Flow cytometry antibodies and clones.

Cells were first incubated with Fc Block (BD Biosciences) (and non-biotinylated human insulin for pancreata) per the manufacturer’s instructions, followed by surface Ab staining (with human insulin for lymph nodes and spleen, mAb123 for pancreata) for 1 hour at 4°C in FACS buffer. Cells were washed three times with FACS buffer and then stained with streptavidin-fluorochrome for 1 hour at 4°C. Cells were then incubated with Foxp3/Transcription Factor Staining buffer (eBioscience, #00-5323-00) for 30 minutes at 4°C. Samples were washed twice with 1x permeabilization buffer from the kit and stained with the intracellular Ab mix in 1x permeabilization buffer overnight at 4°C. After overnight incubation, cells were washed three times with 1x permeabilization buffer, resuspended in FACS buffer, and acquired on the Aurora Spectral Cytometer (Cytek). Cells for single-color controls were prepared similarly as described.

Flow cytometry data were analyzed using FlowJo v10, R v4.5.1, and Cytobank v9.2. The following R packages were used for analyses: dplyr_1.1.4, stringr_1.5.1 readr_2.1.5, magrittr_2.0.3, and cyCombine_0.2.19.

### Minimally Supervised Analysis of Flow Cytometry Data using t-SNE, T-REX, and MEM

FCS files were imported into R and underwent CyCombine normalization ^63^. Batch correction was performed using cyCombine v0.2.19 in R. Data were arcsinh- transformed (cofactor=6000) and down-sampled to the minimum cell count across samples to ensure equal representation. Batch correction was then applied with an 8x8 self-organizing map grid, ’scale’ normalization method, and ’condition’ as covariate to preserve biological differences.

Next, FCS files were concatenated with key labelling conditions (Cre- and Cre+) and imported into Cytobank. Appropriate arcsinh cofactors were set for each marker as previously described ^52^ and exported, and samples were gated on CD45+ B220+ CD19+ live singlet cells. T-REX analysis was performed with appropriate factors set for each population as described, with minimum clustering size set to 25 for Total B cells, 10 for insulin-binding B cells, k value 60-150, and epsilon value 0.3 ^52^. Percent change was calculated based on the abundance of cells from each sample within its KNN region. Identified nodes were merged if multiple identified clusters had identical features. MEM was used (cutoff +2) to provide phenotypic, quantified labels for each identified node ^53^. For pancreatic lymph nodes, B cells underwent initial MEM analysis; GL7, Ki67, CD95, IgM, and CD44 were removed from subsequent analyses to gain depth of MEM labels for less dynamically expressed markers. For pancreata, B cells underwent initial MEM analysis excluding IgG1, as mAb123 is an IgG based antibody which binds to insulin-binding B cells. Following this, GL7, Ki67, IgM, CD23, Igk were removed from subsequently analyses to gain depth of MEM labels for less dynamically expressed markers. T-REX clusters were imported into Cytobank to generate heatmap plots showing markers expression for each cluster. Insulin-binding B cells were additionally analyzed using viSNE on Cytobank with default parameters (iterations=1000, perplexity=30, theta=0.5), with appropriate arcsinh cofactors set for each variable. HINS and IgG1 were omitted from the subsequent marker expression and heatmap analyses for pancreas, and HINS was removed for pancreatic draining lymph nodes. Density plots were created for each concatenated group (split by genotypes, density percent per contour 10%) and used to guide manually drawn clusters. Individual mice expression of each cluster were plotted, and heatmaps were generated to show marker expression for each of these clusters.

### ELISAs

Competitive binding ELISA was used to detect insulin-specific antibodies, as outlined in ^27^. Briefly, 384-well Maxisorp Nunc plates (Thermo Scientific) were coated with 1 μg/ml human insulin in borate-buffered saline overnight at 37°C. Sera were diluted 1:100 in 1X PBS+0.1% Tween-20 (PBS-T) and incubated in wells for 1 hour at room temperature. To determine insulin-specific IgG, parallel samples were incubated in the presence of 100 μg/ml human insulin to enable subtraction of O.D. values from non-inhibited wells. After sera incubation, goat anti-mouse IgG secondary conjugated to alkaline phosphatase (catalogue number 1030-04; Southern Biotech, diluted 1:250) was added. Plates were washed with PBS-T after each of the steps above. Wells were incubated with 10 mg/ml p-nitrophenyl phosphate substrate (Sigma-Aldrich) in a 50 mM potassium carbonate + 1 mM magnesium chloride buffer. Optical density was read at 405 nm after 30 min using a Synergy LX Microplate Autoreader (Bio-Tek).

### Statistical Analyses

Statistical tests are indicated in the corresponding figure legends and significance values were calculated using GraphPad Prism v9.3.1 (GraphPad Software).

### Study approval

All experiments involving animals were approved by Vanderbilt University IACUC.

## Data availability

Supplementary information is available for this manuscript. Values for all data points in graphs are reported in the Supporting Data Sets file.

## Abbreviations

BCR: B cell receptor
EF: extrafollicular
GC: germinal center
NOD: insulin (INS/HINS) non-obese diabetic
SLE: systemic lupus erythematosus
Tfh: T follicular helper
Tfr: T follicular regulatory
Tph: T peripheral helper
T1D: type 1 diabetes

## REFERENCES

1. Evans-Molina C, Oram RA. Teplizumab approval for type 1 diabetes in the USA. The Lancet Diabetes & Endocrinology. 2023;11(2):76–77. doi:10.1016/S2213-8587(22)00390-4

2. Herold KC, Delong T, Perdigoto AL, Biru N, Brusko TM, Walker LSK. The immunology of type 1 diabetes. Nat Rev Immunol. 2024;24(6):435–451. doi:10.1038/s41577-023-00985-4

3. Wherrett DK, Chiang JL, Delamater AM, et al. Defining pathways for development of disease-modifying therapies in children with type 1 diabetes: a consensus report. Diabetes Care. 2015;38(10):1975–1985. doi:10.2337/dc15-1429

4. Pescovitz MD, Greenbaum CJ, Bundy B, et al. B-Lymphocyte Depletion With Rituximab and β-Cell Function: Two-Year Results. Diabetes Care. 2014;37(2):453–459. doi:10.2337/dc13-0626

5. Dörner T, Hansen A. Autoantibodies in normals – the value of predicting rheumatoid arthritis. Arthritis Res Ther. 2004;6(6):282–284. doi:10.1186/ar1456

6. Gómez-Bañuelos E, Fava A, Andrade F. An update on autoantibodies in systemic lupus erythematosus. Curr Opin Rheumatol. 2023;35(2):61–67. doi:10.1097/BOR.0000000000000922

7. Wenzlau JM, Hutton JC. Novel Diabetes Autoantibodies and Prediction of Type 1 Diabetes. Curr Diab Rep. 2013;13(5):10.1007/s11892-013-0405-0409. doi:10.1007/s11892-013-0405-9

8. Achenbach P, Koczwara K, Knopff A, Naserke H, Ziegler AG, Bonifacio E. Mature high- affinity immune responses to (pro)insulin anticipate the autoimmune cascade that leads to type 1 diabetes. J Clin Invest. 2004;114(4):589–597. doi:10.1172/JCI21307

9. Schlosser M, Koczwara K, Kenk H, et al. In insulin-autoantibody-positive children from the general population, antibody affinity identifies those at high and low risk. Diabetologia. 2005;48(9):1830–1832. doi:10.1007/s00125-005-1864-6

10. Yu L, Robles DT, Abiru N, et al. Early expression of antiinsulin autoantibodies of humans and the NOD mouse: Evidence for early determination of subsequent diabetes. Proc Natl Acad Sci U S A. 2000;97(4):1701–1706. doi:10.1073/pnas.040556697

11. Kendall PL, Case JB, Sullivan AM, et al. Tolerant anti-insulin B cells are effective APCs. J Immunol. 2013;190(6):2519–2526. doi:10.4049/jimmunol.1202104

12. Crotty S. T Follicular Helper Cell Biology: A Decade of Discovery and Diseases. Immunity. 2019;50(5):1132–1148. doi:10.1016/j.immuni.2019.04.011

13. Gd V, Mc N. Germinal Centers. Annual review of immunology. 2022;40. doi:10.1146/annurev-immunol-120419-022408

14. Edner NM, Heuts F, Thomas N, et al. Follicular helper T cell profiles predict response to costimulation blockade in type 1 diabetes. Nat Immunol. 2020;21(10):1244–1255. doi:10.1038/s41590-020-0744-z

15. Gearty SV, Dündar F, Zumbo P, et al. An autoimmune stem-like CD8 T cell population drives type 1 diabetes. Nature. 2022;602(7895):156–161. doi:10.1038/s41586-021-04248-x

16. Russell WE, Bundy BN, Anderson MS, et al. Abatacept for Delay of Type 1 Diabetes Progression in Stage 1 Relatives at Risk: A Randomized, Double-Masked, Controlled Trial. Diabetes Care. 2023;46(5):1005–1013. doi:10.2337/dc22-2200

17. Viisanen T, Ihantola EL, Näntö-Salonen K, et al. Circulating CXCR5+PD-1+ICOS+ Follicular T Helper Cells Are Increased Close to the Diagnosis of Type 1 Diabetes in Children With Multiple Autoantibodies. Diabetes. 2017;66(2):437–447. doi:10.2337/db16-0714

18. Yang K, Zhang Y, Ding J, Li Z, Zhang H, Zou F. Autoimmune CD8+ T cells in type 1 diabetes: from single-cell RNA sequencing to T-cell receptor redirection. Front Endocrinol (Lausanne). 2024;15:1377322. doi:10.3389/fendo.2024.1377322

19. Wan X, Thomas JW, Unanue ER. Class-switched anti-insulin antibodies originate from unconventional antigen presentation in multiple lymphoid sites. J Exp Med. 2016;213(6):967–978. doi:10.1084/jem.20151869

20. Nakayama M, Michels AW. Using the T Cell Receptor as a Biomarker in Type 1 Diabetes. Front Immunol. 2021;12:777788. doi:10.3389/fimmu.2021.777788

21. Bonami RH, Nyhoff LE, McNitt DH, et al. T-B Lymphocyte Interactions Promote Type 1 Diabetes Independently of SLAM-Associated Protein. J Immunol. 2020;205(12):3263–3276. doi:10.4049/jimmunol.1900464

22. Wellford SA, Schwartzberg PL. Help me help you: emerging concepts in T follicular helper cell differentiation, identity, and function. Curr Opin Immunol. 2024;87:102421. doi:10.1016/j.coi.2024.102421

23. Basso K, Dalla-Favera R. Roles of BCL6 in normal and transformed germinal center B cells. Immunological Reviews. 2012;247(1):172–183. doi:10.1111/j.1600-065X.2012.01112.x

24. Choi J, Diao H, Faliti CE, et al. Bcl-6 is the nexus transcription factor of T follicular helper cells via repressor-of-repressor circuits. Nat Immunol. 2020;21(7):777–789. doi:10.1038/s41590-020-0706-5

25. Choi J, Crotty S. Bcl6-Mediated Transcriptional Regulation of Follicular Helper T cells (TFH). Trends Immunol. 2021;42(4):336–349. doi:10.1016/j.it.2021.02.002

26. Hatzi K, Melnick A. Breaking bad in the germinal center: how deregulation of BCL6 contributes to lymphomagenesis. Trends Mol Med. 2014;20(6):343–352. doi:10.1016/j.molmed.2014.03.001

27. McNitt DH, Williams JM, Santitoro JG, Kim J, Thomas JW, Bonami RH. Type 1 Diabetes Depends on CD4-Driven Expression of the Transcriptional Repressor Bcl6. Diabetes. 2025;74(6):921–932. doi:10.2337/db23-0709

28. Hollister K, Kusam S, Wu H, et al. Insights into the Role of Bcl6 in Follicular Helper T Cells Using a New Conditional Mutant Mouse Model. J Immunol. 2013;191(7):3705–3711. doi:10.4049/jimmunol.1300378

29. Vremec D, Pooley J, Hochrein H, Wu L, Shortman K. CD4 and CD8 expression by dendritic cell subtypes in mouse thymus and spleen. J Immunol. 2000;164(6):2978–2986. doi:10.4049/jimmunol.164.6.2978

30. Henry RA, Kendall PL, Thomas JW. Autoantigen-Specific B-Cell Depletion Overcomes Failed Immune Tolerance in Type 1 Diabetes. Diabetes. 2012;61(8):2037–2044. doi:10.2337/db11-1746

31. Akama-Garren EH, Carroll MC. T Cell Help in the Autoreactive Germinal Center. Scand J Immunol. 2022;95(6):e13192. doi:10.1111/sji.13192

32. Olivieri G, Cotugno N, Palma P. Emerging insights into atypical B cells in pediatric chronic infectious diseases and immune system disorders: T(o)-bet on control of B-cell immune activation. Journal of Allergy and Clinical Immunology. 2024;153(1):12–27. doi:10.1016/j.jaci.2023.10.009

33. Stebegg M, Kumar SD, Silva-Cayetano A, Fonseca VR, Linterman MA, Graca L. Regulation of the Germinal Center Response. Front Immunol. 2018;9. doi:10.3389/fimmu.2018.02469

34. Jenks SA, Cashman KS, Zumaquero E, et al. Distinct Effector B Cells Induced by Unregulated Toll-like Receptor 7 Contribute to Pathogenic Responses in Systemic Lupus Erythematosus. Immunity. 2018;49(4):725–739.e6. doi:10.1016/j.immuni.2018.08.015

35. Liu X, Yan X, Zhong B, et al. Bcl6 expression specifies the T follicular helper cell program in vivo. J Exp Med. 2012;209(10):1841–1852, S1-24. doi:10.1084/jem.20120219

36. Nickerson KM, Smita S, Hoehn KB, et al. Age-associated B cells are heterogeneous and dynamic drivers of autoimmunity in mice. J Exp Med. 2023;220(5):e20221346. doi:10.1084/jem.20221346

37. Song W, Antao OQ, Condiff E, et al. Development of Tbet- and CD11c-expressing B cells in a viral infection requires T follicular helper cells outside of germinal centers. Immunity. 2022;55(2):290–307.e5. doi:10.1016/j.immuni.2022.01.002

38. Wang S, Wang J, Kumar V, et al. IL-21 drives expansion and plasma cell differentiation of autoreactive CD11chiT-bet+ B cells in SLE. Nat Commun. 2018;9(1):1758. doi:10.1038/s41467-018-03750-7

39. Sachinidis A, Trachana M, Taparkou A, et al. Characterization of T-bet expressing B cells in lupus patients indicates a putative prognostic and therapeutic value of these cells for the disease. Clin Exp Immunol. 2025;219(1):uxaf008. doi:10.1093/cei/uxaf008

40. Wu C, Fu Q, Guo Q, et al. Lupus-associated atypical memory B cells are mTORC1- hyperactivated and functionally dysregulated. Ann Rheum Dis. 2019;78(8):1090–1100. doi:10.1136/annrheumdis-2019-215039

41. Felton JL, Maseda D, Bonami RH, Hulbert C, Thomas JW. Anti-Insulin B Cells Are Poised for Antigen Presentation in Type 1 Diabetes. J Immunol. 2018;201(3):861–873. doi:10.4049/jimmunol.1701717

42. Hulbert C, Riseili B, Rojas M, Thomas JW. B cell specificity contributes to the outcome of diabetes in nonobese diabetic mice. J Immunol. 2001;167(10):5535–5538. doi:10.4049/jimmunol.167.10.5535

43. Williams JM, Bonami RH, Hulbert C, Thomas JW. Reversing Tolerance in Isotype Switch- Competent Anti-Insulin B Lymphocytes. J Immunol. 2015;195(3):853–864. doi:10.4049/jimmunol.1403114

44. Lu Y, Craft J. T Follicular Regulatory Cells: Choreographers of Productive Germinal Center Responses. Front Immunol. 2021;12:679909. doi:10.3389/fimmu.2021.679909

45. Kendall PL, Yu G, Woodward EJ, Thomas JW. Tertiary lymphoid structures in the pancreas promote selection of B lymphocytes in autoimmune diabetes. J Immunol. 2007;178(9):5643–5651. doi:10.4049/jimmunol.178.9.5643

46. Hamel KM, Liarski, Vladimir M., and Clark MR. Germinal Center B-cells. Autoimmunity. 2012;45(5):333-347. doi:10.3109/08916934.2012.665524

47. Duy C, Yu JJ, Nahar R, et al. BCL6 is critical for the development of a diverse primary B cell repertoire. Journal of Experimental Medicine. 2010;207(6):1209–1221. doi:10.1084/jem.20091299

48. Kitayama D, Sakamoto A, Arima M, Hatano M, Miyazaki M, Tokuhisa T. A role for Bcl6 in sequential class switch recombination to IgE in B cells stimulated with IL-4 and IL-21. Mol Immunol. 2008;45(5):1337–1345. doi:10.1016/j.molimm.2007.09.007

49. Roco JA, Mesin L, Binder SC, et al. Class-Switch Recombination Occurs Infrequently in Germinal Centers. Immunity. 2019;51(2):337–350.e7. doi:10.1016/j.immuni.2019.07.001

50. Koczwara K, Schenker M, Schmid S, Kredel K, Ziegler AG, Bonifacio E. Characterization of antibody responses to endogenous and exogenous antigen in the nonobese diabetic mouse. Clin Immunol. 2003;106(2):155–162. doi:10.1016/s1521-6616(02)00040-2

51. McNitt DH, Joosse BA, Thomas JW, Bonami RH. Productive Germinal Center Responses Depend on the Nature of Stimuli Received by Anti-Insulin B Cells in Type 1 Diabetes– Prone Mice. Immunohorizons. 2023;7(6):384–397. doi:10.4049/immunohorizons.2300036

52. Barone SM, Paul AG, Muehling LM, et al. Unsupervised machine learning reveals key immune cell subsets in COVID-19, rhinovirus infection, and cancer therapy. Elife. 2021;10:e64653. doi:10.7554/eLife.64653

53. Diggins KE, Greenplate AR, Leelatian N, Wogsland CE, Irish JM. Characterizing cell subsets using marker enrichment modeling. Nat Methods. 2017;14(3):275–278. doi:10.1038/nmeth.4149

54. Kramer KJ, Wilfong EM, Voss K, et al. Single-cell profiling of the antigen-specific response to BNT162b2 SARS-CoV-2 RNA vaccine. Nat Commun. 2022;13(1):3466. doi:10.1038/s41467-022-31142-5

55. Groocock L, Deb G, Zhu J, et al. BMS-986458 a Potential First-in-Class, Highly Selective, Potent and Well Tolerated BCL6 Ligand Directed Degrader (LDD) Demonstrates Multi- Modal Anti-Tumor Efficacy for the Treatment of B-Cell Non-Hodgkin’s Lymphoma. Blood. 2024;144(Supplement 1):957. doi:10.1182/blood-2024-210951

56. Henry RA, Kendall PL. CXCL13 blockade disrupts B lymphocyte organization in tertiary lymphoid structures without altering B cell receptor bias or preventing diabetes in nonobese diabetic mice. J Immunol. 2010;185(3):1460–1465. doi:10.4049/jimmunol.0903710

57. Ambegaonkar AA, Holla P, Dizon BL, Sohn H, Pierce SK. Atypical B cells in chronic infectious diseases and systemic autoimmunity: puzzles with many missing pieces. Curr Opin Immunol. 2022;77:102227. doi:10.1016/j.coi.2022.102227

58. Andrews LP, Vignali KM, Szymczak-Workman AL, et al. A Cre-driven allele-conditioning line to interrogate CD4+ conventional T cells. Immunity. 2021;54(10):2209–2217.e6. doi:10.1016/j.immuni.2021.08.029

59. Zuccarino-Catania GV, Sadanand S, Weisel FJ, et al. CD80 and PD-L2 define functionally distinct memory B cell subsets that are independent of antibody isotype. Nat Immunol. 2014;15(7):631–637. doi:10.1038/ni.2914

60. Markle JGM, Frank DN, Mortin-Toth S, et al. Sex Differences in the Gut Microbiome Drive Hormone-Dependent Regulation of Autoimmunity. Science. 2013;339(6123):1084-1088. doi:10.1126/science.1233521

61. Kendall PL, Moore DJ, Hulbert C, Hoek KL, Khan WN, Thomas JW. Reduced diabetes in btk-deficient nonobese diabetic mice and restoration of diabetes with provision of an anti-insulin IgH chain transgene. J Immunol. 2009;183(10):6403–6412. doi:10.4049/jimmunol.0900367

62. Henry-Bonami RA, Williams JM, Rachakonda AB, Karamali M, Kendall PL, Thomas JW. B Lymphocyte “Original Sin” in the Bone Marrow Enhances Islet Autoreactivity in Type 1 Diabetes-Prone NOD mice,,. J Immunol. 2013;190(12):5992–6003. doi:10.4049/jimmunol.1201359

63. Pedersen CB, Dam SH, Barnkob MB, et al. cyCombine allows for robust integration of single-cell cytometry datasets within and across technologies. Nat Commun. 2022;13(1):1698. doi:10.1038/s41467-022-29383-5

