## Supplemental Figures for "BCL6 in T cells promotes type 1 diabetes by redirecting fates of insulin-autoreactive B lymphocytes"

**SUPPLEMENTAL MATERIALS**

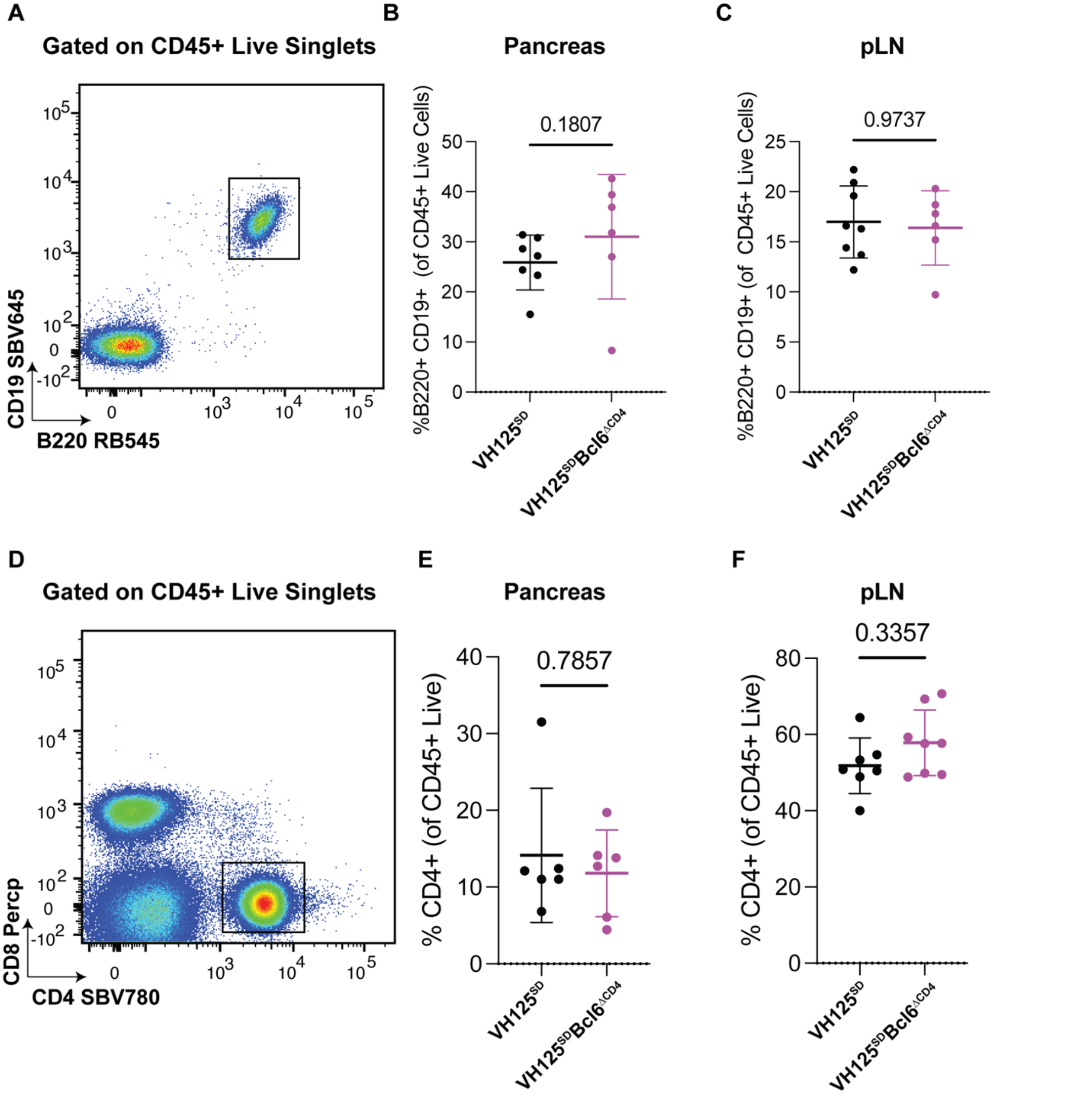


**Supplemental Figure 1: Total B and CD4+ T cells are not significantly altered in the pancreas and pancreatic lymph nodes by *Cd4*-Cre loss of *Bcl6*.** Cells were isolated from spleen and pancreas from 8-12-week-old VH125$^{SD}$.NOD and VH125$^{SD}$.Bcl6$^{\Delta CD4}$.NOD mice. **(A)** Representative flow plots defining B cells as live singlet B220$^{+}$ CD19$^{+}$ CD45$^{+}$ lymphocytes. **(B-C)** % B cells identified as in *(A)* is shown for **(B)** pancreas and **(C)** pancreatic lymph nodes. **(D)** Representative flow plots defining CD4+ T cells as live singlet CD4$^{+}$ CD45$^{+}$ lymphocytes. **(E-F)** % CD4+ T cells identified as in *(D)* is shown for **(E)** pancreas and **(F)** pancreatic lymph nodes. n = 6-8 mice per group, significance determined by Mann-Whitney U-test.

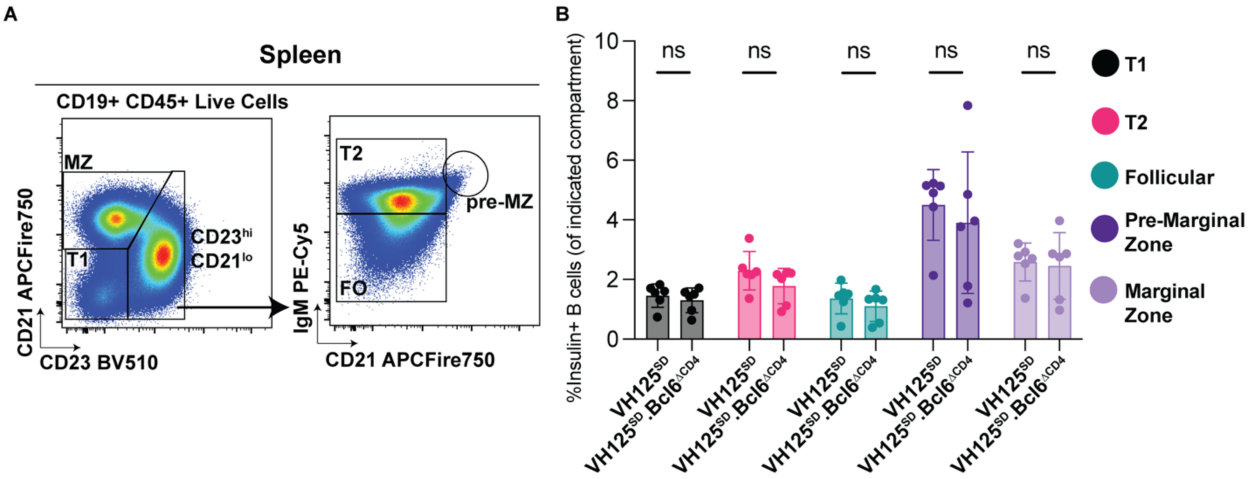


**Supplemental Figure 2: B cell development of insulin-binding B cells are not reduced by *Cd4*-Cre-driven *Bcl6* loss.** Cells were isolated from spleen and pancreas from 8-12-week-old VH125$^{SD}$.NOD and VH125$^{SD}$.Bcl6$^{\Delta CD4}$.NOD mice. **(A)** Following gating of CD19+ CD45+ Live cells, splenic B cell subsets were identified as follows: T1 (CD21$^{low}$ CD23$^{low}$ IgM$^{high}$), T2 (CD21$^{low}$ CD23$^{high}$ IgM$^{high}$), FO (CD21$^{low}$ CD23$^{high}$ IgM$^{low}$), Pre-MZ (CD21$^{high}$ CD23$^{high}$ IgM$^{high}$) and MZ (CD21$^{high}$ CD23$^{low}$ IgM$^{high}$), as shown in representative flow plots. **(B)** Within each B cell subset, the percentage of insulin-binding B cells was identified within each B cell subset. n=6-8 mice per group, Kruskal-Wallis non-parametric statistical tests with post-hoc uncorrected Dunn's test were used, ns = not significant. Bars indicate mean +/- SD.

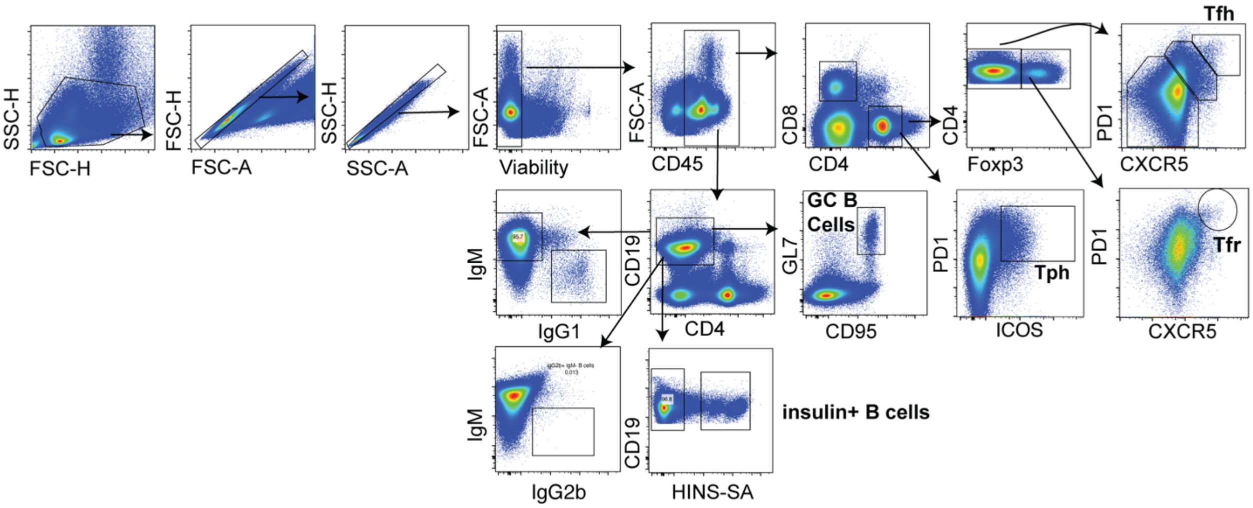


**Supplemental Figure 3: Gating strategy to identify CD4+, CD8+, and CD19+ populations.** Spleens were isolated from VH125$^{SD}$.NOD and flow cytometry staining and gating was performed as shown in representative plots. Similar gating strategies were performed for other lymph nodes (pancreatic and mesenteric lymph nodes) and pancreata.

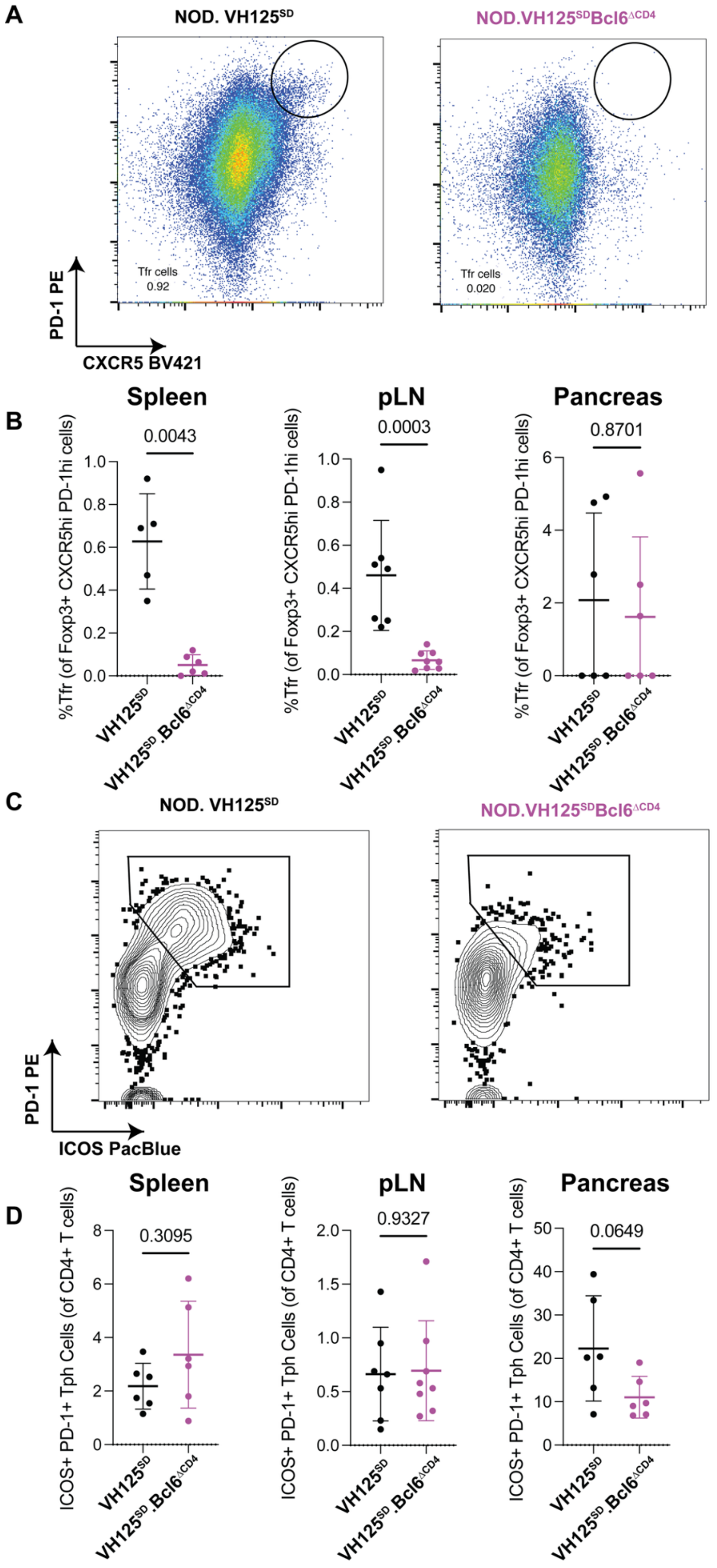
A
NOD. VH125SD
NOD.VH125SDBcl6ΔCD4
Tfr cells
0.92
Tfr cells
0.020
PD-1 PE
CXCR5 BV421
B
Spleen
pLN
Pancreas
0.0043
0.0003
0.8701
%Tfr (of Foxp3+ CXCR5hi PD-1hi cells)
VH125SD
VH125SD.Bcl6ΔCD4
C
NOD. VH125SD
NOD.VH125SDBcl6ΔCD4
PD-1 PE
ICOS PacBlue
D
Spleen
pLN
Pancreas
0.3095
0.9327
0.0649
ICOS+ PD-1+ Tph Cells (of CD4+ T cells)
VH125SD
VH125SD.Bcl6ΔCD4

**Supplemental Figure 4: Loss of *Bcl6* reduces T follicular regulatory and T peripheral helper populations in spleen and pancreas.** Cells from spleen, pancreatic lymph nodes (pLNs), and pancreas were isolated from 8-12-week-old VH125$^{SD}$.NOD and VH125$^{SD}$.Bcl6$^{\Delta CD4}$.NOD mice. **(A)** Representative flow cytometry plots from spleen identify T follicular regulatory (Tfr) cells (live singlet CD45$^{+}$ CD4$^{+}$ Foxp3$^{+}$ CXCR5$^{hi}$ PD-1$^{hi}$lymphocytes), significance determined by Mann-Whitney U-test. **(B)** % Tfr cells in spleen, pLNs, and pancreas is plotted for individual mice. **(C)** Representative flow cytometry plots identify pancreatic T peripheral helper (Tph) cells in spleen live singlet CD45$^{+}$ CD4$^{+}$ ICOS$^{+}$ PD-1$^{+}$ lymphocytes). **(D)** % Tph cells in spleen, pLNs, and pancreas is plotted for individual mice. n=6 mice per group, significance determined by Mann-Whitney U-test.

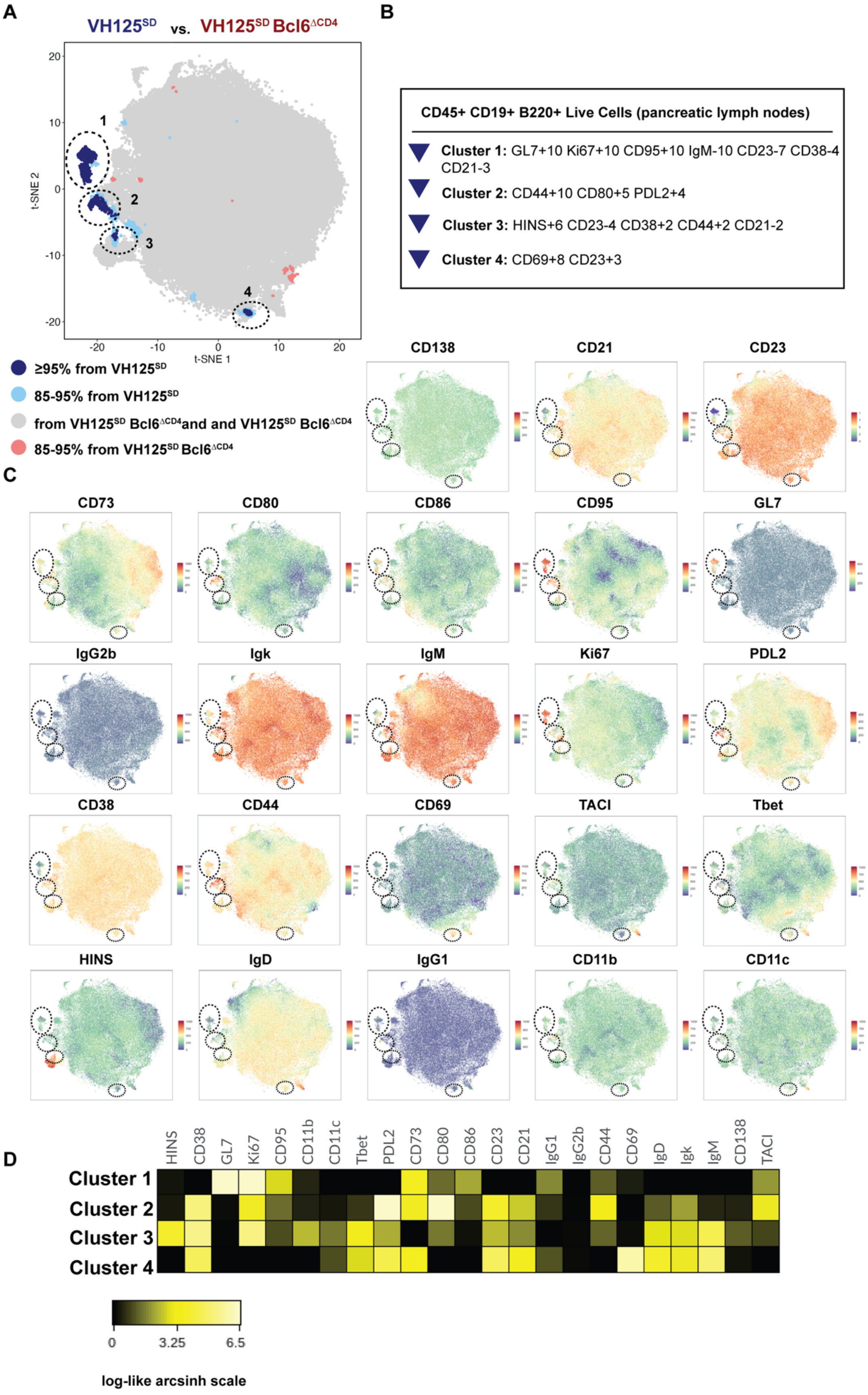
A
VH125SD vs. VH125SD Bcl6ΔCD4
t-SNE 2
t-SNE 1
1
2
3
4
≥95% from VH125SD
85-95% from VH125SD
from VH125SD Bcl6ΔCD4 and and VH125SD Bcl6ΔCD4
85-95% from VH125SD Bcl6ΔCD4
B
CD45+ CD19+ B220+ Live Cells (pancreatic lymph nodes)
Cluster 1: GL7+10 Ki67+10 CD95+10 IgM-10 CD23-7 CD38-4 CD21-3
Cluster 2: CD44+10 CD80+5 PDL2+4
Cluster 3: HINS+6 CD23-4 CD38+2 CD44+2 CD21-2
Cluster 4: CD69+8 CD23+3
C
CD138
CD21
CD23
CD73
CD80
CD86
CD95
GL7
IgG2b
Igk
IgM
Ki67
PDL2
CD38
CD44
CD69
TACI
Tbet
HINS
IgD
IgG1
CD11b
CD11c
D
HINS
CD38
GL7
Ki67
CD95
CD11b
CD11c
Tbet
PDL2
CD73
CD80
CD86
CD23
CD21
IgG1
IgG2b
CD44
CD69
IgD
Igk
IgM
CD138
TACI
Cluster 1
Cluster 2
Cluster 3
Cluster 4
0
3.25
6.5
log-like arcsinh scale

**Supplemental Figure 5: Loss of *Bcl6* via *Cd4*-Cre leads to contracted populations of B cells with phenotypic attributes of GC, activated, and atypical memory B cells in VH125$^{SD}$.NOD mice.** Pancreatic draining lymph nodes were harvested from pre-diabetic, 8-12 week-old VH125$^{SD}$.NOD mice. B220+ CD19+ CD45+ live cells from n = 12-14 mice were normalized via CyCombine and concatenated into two groups, *Bcl6*-sufficient (VH125$^{SD}$) and *Bcl6*-deficient (VH125$^{SD}$Bcl6$^{\Delta CD4}$.NOD). **(A)** t-SNE was used to perform dimensionality reduction based on phenotypic marker expression profiles. Following this, the minimally supervised analysis tool, Tracking Responders Expanding (T-REX) was used to identify populations that were increased (red: > 95% or peach: 85-95% confidence levels) or decreased (blue: > 95% or light blue: 85-95% confidence levels) by *Cd4*-Cre loss of *Bcl6* in this model, as illustrated in the t-SNE plot. **(B)** Marker Enrichment Modeling (MEM) labels show the phenotypic marker features that define each of the contracted populations. Features enriched by at least +2 or -2 on a scale from 0 to 10 are shown. **(C)** A rainbow intensity scale indicates expression levels of the phenotypic markers indicated at the top of each t-SNE plot with red representing high levels and blue representing low levels of expression. CD45, B220, CD19, and the viability dye expression was omitted from the t-SNE analysis given these markers were used to gate on the parent population that went into phenotypic clustering via t-SNE and T-REX visualization and analysis. **(D)** Heatmap shows relative expression of the indicated markers for the contracted clusters identified by T-REX in *(A)*. Areas of high expression (yellow) and low expression (black) are shown via a log-like arcsinh scale. Data were scaled and represented based on individual marker expression.

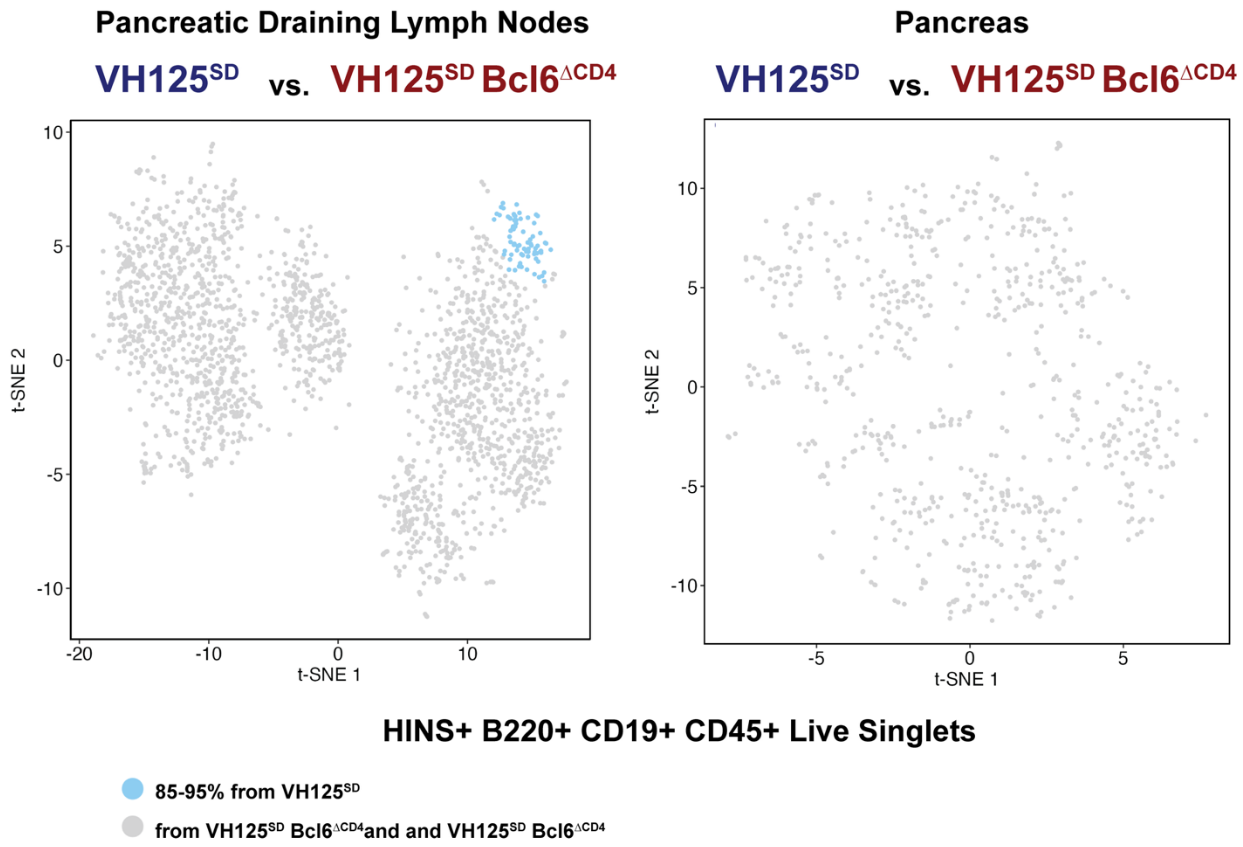


**Supplemental Figure 6: T-REX fails to identify key anti-insulin B cell changes in pancreatic draining lymph nodes and pancreata.** Following cyCombine normalization analysis, anti-insulin B cells from pancreatic draining lymph nodes in both $VH125^{SD}$.NOD and $VH125^{SD}$.$Bcl6^{\Delta CD4}$.NOD mice underwent T-REX analysis. Areas of contracted B cell populations with *Bcl6* loss are shown in blue. Minimum number of required events for MEM Label=10, K=150, epsilon=0.3, Perplexity=100, Iterations=1000.

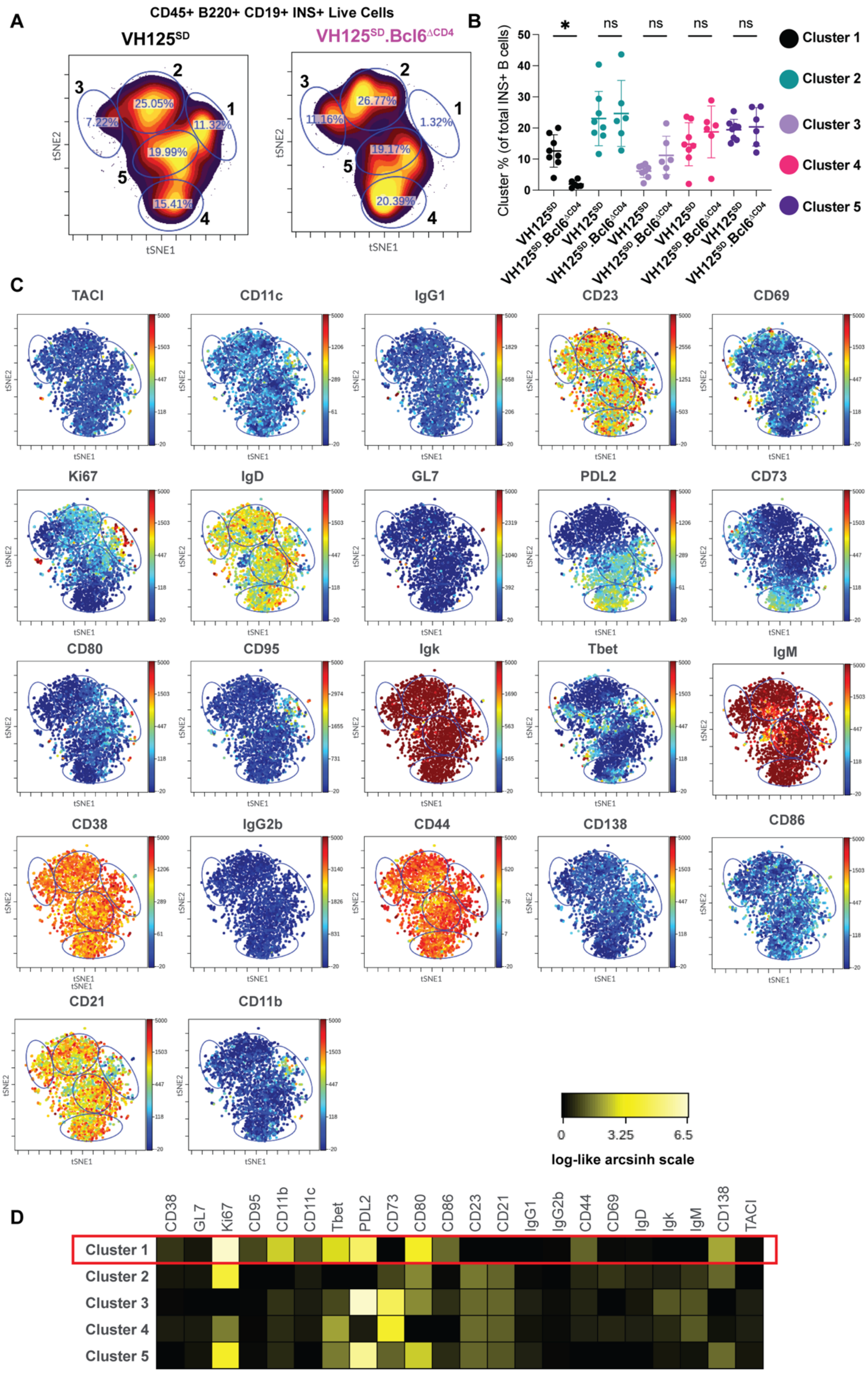
A
CD45+ B220+ CD19+ INS+ Live Cells
VH125SD
VH125SD.Bcl6ΔCD4
25.05%
11.32%
7.22%
19.99%
15.41%
26.77%
11.16%
1.32%
19.17%
20.39%
tSNE1
tSNE2
B
Cluster % (of total INS+ B cells)
*
ns
ns
ns
ns
Cluster 1
Cluster 2
Cluster 3
Cluster 4
Cluster 5
C
TACI
CD11c
IgG1
CD23
CD69
Ki67
IgD
GL7
PDL2
CD73
CD80
CD95
Igk
Tbet
IgM
CD38
IgG2b
CD44
CD138
CD86
CD21
CD11b
0
3.25
6.5
log-like arcsinh scale
D
CD38
GL7
Ki67
CD95
CD11b
CD11c
Tbet
PDL2
CD73
CD80
CD86
CD23
CD21
IgG1
IgG2b
CD44
CD69
IgD
Igk
IgM
CD138
TACI
Cluster 1
Cluster 2
Cluster 3
Cluster 4
Cluster 5

**Supplemental Figure 7: Atypical memory-like B cell populations are decreased among insulin-binding B cells with *Cd4*-Cre loss of *Bcl6* in VH125$^{SD}$.NOD mice in pancreatic draining lymph nodes.** Pancreatic draining lymph nodes were harvested from pre-diabetic, 8–12-week-old VH125$^{SD}$.NOD mice. Insulin-binding B220+ CD19+ CD45+ live singlets cells (n = 6-7 mice per genotype) were normalized via CyCombine and concatenated into two groups, *Bcl6*-sufficient (VH125$^{SD}$) and *Bcl6*-deficient (VH125$^{SD}$Bcl6$^{\Delta CD4}$.NOD). **(A)** t-SNE was used to perform dimensionality reduction based on phenotypic marker expression profiles. Clusters were manually defined as indicated. **(B)** The cluster frequency is shown for each genotype, with individual mice plotted and means indicated. * $p < 0.05$, Kruskal-Wallis test with post-hoc uncorrected Dunn's test. **(C)** A rainbow intensity scale indicates expression levels of the phenotypic markers indicated at the top of each t-SNE plot. Insulin-binding, CD45, B220, CD19, and viability dye markers were omitted from the t-SNE analysis given they were used in parent population gating upstream of t-SNE analysis. **(D)** Heatmap shows relative expression of the indicated markers for each cluster defined as in *(A)* using a log-like arcsinh scale and appropriate arcsinh factors.
